# Composite branch construction by dual autozooidal budding modes in hornerids (Bryozoa: Cyclostomatida)

**DOI:** 10.1101/2022.01.30.478354

**Authors:** P. B. Batson, Y. Tamberg, P.D. Taylor

## Abstract

Horneridae (Cyclostomata; Cancellata) is a family of marine bryozoans that form tree-like colonies bearing functionally unilaminate branches. Colony development in this clade is not well understood. We used micro-CT and SEM to trace zooidal budding in *Hornera* from the ancestrula onwards. Results show that hornerid branches are constructed by dual zooidal budding modes occurring synchronously at two separate budding sites at the growing tips. Frontal autozooids bud from a multizooidal budding lamina. Lateral autozooids bud from discrete abfrontal budding loci by ‘exomural budding’, a previously undescribed form of budding centred on hypostegal pores in interzooidal grooves on the colonial body wall. These two budding modes are integrated during primary branch morphogenesis, forming composite, developmentally bilaminate, branches. Tracing development back to the founding zooids shows that the ancestrula and adventitious periancestrular autozooids are the first-formed lateral autozooids. They grow as a vertical stem onto which new laterals bud exomurally, intercalating with earlier autozooids to form an expanding funnel. The inner wall of this funnel becomes a ring-shaped budding lamina, from which the first frontal autozooids bud centripetally. When the branch crown splits, the ring lamina is subdivided, each fragment becoming a paramedial budding lamina in its own branch, while new laterals continue to be budded exomurally onto the frontal walls of existing laterals. Co-occurrence of two rare cyclostome frontal budding types—adventitious budding on the ancestrula and exomural budding on the branches—suggests they may be homologous. Patterns of exomural budding vary among hornerids. Narrow-branched taxa possess a single medial line of budding sites on the abfrontal wall; wide-branched species have up to six parallel lines of loci. Exomural budding also occurs sporadically in *Calvetia osheai*, a radial-branching hornerid. Future studies of cancellate taxonomy and phylogeny may benefit from morphological concepts presented here.

## 1. Introduction

Bryozoa is a phylum of colonial, mostly sessile, aquatic invertebrates composed of physically and physiologically connected zooids (Ryland, 1970). Each zooid is budded asexually and is homologous with a unitary organism. In most marine taxa zooids are small (< 1 mm), tubular or box-shaped, and are often calcified (Schwaha et al., 2020). Polymorphism is widespread, with zooids being specialised for different functions within the colony—e.g., feeding (autozooids), structural roles (kenozooids) and reproduction (e.g. gonozooids, ovicells) (Schack et al., 2018).

Bryozoans are suspension feeders. Each autozooid contains a polypide, which captures microscopic food particles using a protrusible ring of ciliated feeding tentacles (Winston, 1978). The polypide bauplan is conserved at the ordinal level (e.g. Boardman & McKinney, 1985; Tamberg et al., 2021). In contrast, the arrangements of zooidal modules within a bryozoan colony vary greatly within families, genera and species, resulting in a diverse array of ‘second-order’ morphologies (Hageman, 2003). Location, timing and mode of zooidal budding events are key determinants of the zooid arrangement (Lidgard, 1985). In this article we describe a previously unknown mode of zooidal budding and colony construction in a family of cyclostome bryozoans.

### 1.1. Colony development in Horneridae

Horneridae Smitt, 1867 is a cosmopolitan family of cyclostome bryozoans (class Stenolaemata, suborder Cancellata). It is exemplified by *Hornera* Lamouroux, 1821, species of which develop erect, branching colonies up to ~ 150 mm in height. A typical colony has a wide basal attachment disc supporting a short basal stem and a multi-branched crown giving rise to the main branches. Fan-like, tree-like or fenestrate *Hornera* species occur on continental shelves and slopes, where they can be locally abundant and/or ecologically important (e.g. Batson & Probert, 2000; Wood et al., 2012; Harmelin, 2020). Branches are functionally unilaminate, with apertures of the tubular feeding autozooids opening only on their front sides (Figure 1A). The number of autozooidal chambers in a branch cross-section varies among taxa, from ~4 to 60. Horneridae is moderately diverse, with ~ 100 described living and fossil species, ranging from the Eocene to the present-day (Mongereau, 1972; Smith et al., 2008; Harmelin, 2020).

**Figure 1.**
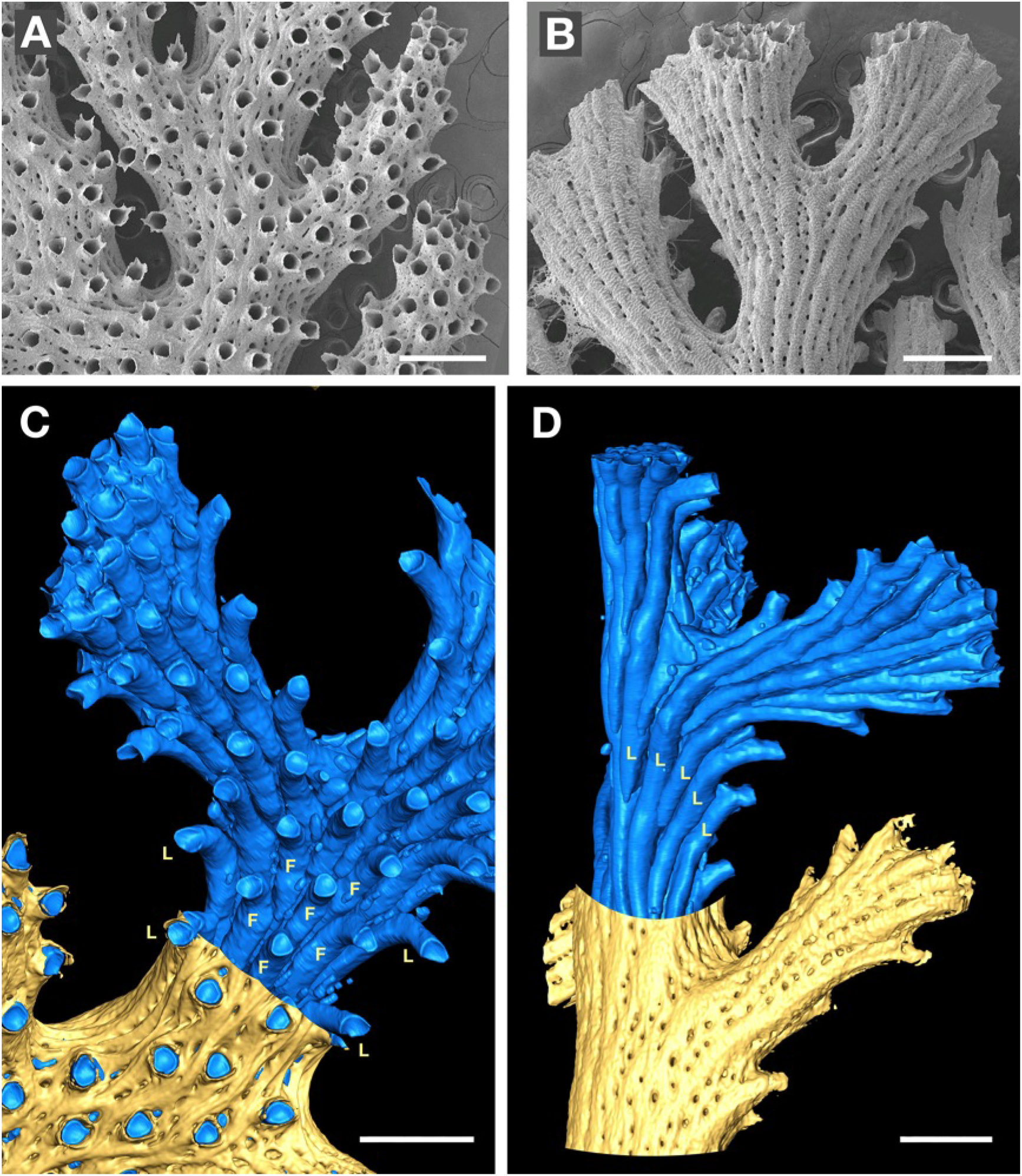
Overview of hornerid skeletal morphology. **A.** SEM of frontal surface of a *Hornera* sp. 1. branch bearing autozooidal apertures with dentate peristomes; smaller openings of cancelli also visible. **B.** Abfrontal branch surface from the same colony; lines of cancelli, pustulose secondary calcification extends to the branch tips. **C.** Micro-CT interior/exterior reconstruction of frontal surface of *Hornera* sp. 1; most of the fine tubes (cancelli/kenozooids) have been removed during model processing. Autozooidal chambers (blue) are surrounded by secondarily calcified body wall (yellow). Frontal autozooids (F) bordered by frontolaterally curved lateral autozooids (L). **D.** Abfrontal surface of same branch bearing only lateral autozooids (L). Scalebars: A, B, 500 μm; C,D, ~500 μm.

Hornerids are among the most-heavily calcified living cyclostomes (Borg, 1926; Batson et al., 2021). Secondary branch thickening begins immediately behind the branch growing tip, which is itself formed by distal extension of the tubular autozooid chambers (Taylor & Jones, 1993). The thick ‘extrazooidal’ skeleton is characterised by longitudinal ridges (nervi) and grooves (sulci), and is penetrated by fine tubules (cancelli) (Figure 1A, B) connecting the outer hypostegal cavity to the zooidal chambers via hypostegal pores. Extrazooidal skeleton is secreted onto the exterior of the autozooidal bundle by an outer, colony-enclosing epithelium (Borg, 1926; Boardman, 1998). This epithelium is part of the interior frontal wall, a body wall type present in most cyclostome suborders (Borg, 1926; Taylor, 2000) and in the related, extinct, palaeostomate bryozoans (Borg, 1965; Ma et al., 2014).

In hornerids the interior frontal wall appears to originate very early in development of the ancestrula (the colony-founding zooid) compared to equivalent walls in other interior-walled cyclostome clades (Batson et al., 2019). Early development of this wall enables periancestrular autozooids to bud adventitiously from (and onto) the broad roof of the ancestrula within 1–2 days of first calcification (Batson et al. 2019).

Formation of the interior frontal wall, the early astogeny of the colony, and subsequent patterns of zooidal budding appear to be linked in *Hornera*, although in ways that are not well understood (e.g., Borg, 1926, p. 305). This relationship has the potential to influence branch construction and the development of secondary wall calcification—two traits that make hornerids so distinctive among cyclostomes (e.g. Borg, 1926; Boardman, 1998; Harmelin, 2020). Unfortunately, study of zooidal budding and colony development in hornerid cyclostomes has long been impeded by their thick secondary wall calcification.

Hornerids have two morphologically distinct types of autozooids: frontals and laterals (Figure 1A–D). This difference was first recognised by Boardman (1998, figures 26–28), who referred to them as ‘polymorphic feeding zooids’ based on their relative size, shape and position. New chambers of both zooid types arise at growing tips of branches. Frontal autozooids are concentrated on the frontal side of branches. Initially, their chambers grow distally within a well-developed zooidal endozone; later in development each frontal zooid separates from those around it and bends frontally, its distal portions opening onto the frontal surface of the branch (Figure 1A, C).

Lateral autozooids comprise a separate, unbroken layer beneath the frontal zooids, closely ‘cupping’ the frontals on their abfrontal and lateral sides (Figure 1D). As the lateral autozooids grow distally, their chambers also ‘migrate’ around the perimeter of the endozone, eventually opening fronto-laterally onto the colony surface in alternating series (Figure 1C). This growth pattern often results in a distinct ‘herringbone’ arrangement of laterally divergent zooids centred along the midline of the abfrontal branch wall (Figure 1D; see also Boardman, 1998, figure 26B).

Despite their close proximity, frontal and lateral zooids remain separate, usually forming a distinct bilayer throughout development. As the zooids lengthen and turn away from the endozone, the space between diverging chambers soon becomes occupied by secondary calcification, leaving only the zooidal peristomes visible proximal of the growing tip. This tendency, combined with the complex 3D arrangement of zooidal chambers, makes it difficult to trace the origins of individual autozooids.

How and where do the hornerid autozooids arise? Various interpretations of zooidal budding in *Hornera* have been offered. Hennig (1911, p. 37) reported “zooecial ducts [autozooids] emerging from an elongated tube inside the cortical part of the dorsal [abfrontal] side and extending from there obliquely anteriorly and upwards towards the oral side” (translated from the French original). Canu & Bassler (1920, p. 796) disagreed with Hennig, adding: “The successive ramification of the tubes is identical with that of other families.” Currently, the prevailing model of zooidal budding is that of Borg (1926; p. 306), who wrote that all autozooids in *H. antarctica* are budded “at the basal wall of the zoarium”, with the distal parts of zooids “gradually forced away from the basal side of the stem” [by budding of new zooids]. He concurred with Smitt’s (1867, p. 471) view that budding is “as a rule confined to the median part of the stem”. Borg later included budding from the basal wall as a diagnostic character in his redescription of the Horneridae (1944, p. 185).

Since Borg (1944), most studies give descriptions compatible with the presence of a well-defined basal budding lamina. Schäfer (1991, figure 56) illustrated a longitudinal section of a *Hornera frondiculata* branch consistent with Borg’s interpretation of zooidal budding in this genus, as did Brood (1976). However, Drexler (1976; p. 20) described budding as more variable in some late Eocene hornerid fossils, in which “budding was concentrated at the branch distal growing tip (“common bud”) but either occurred throughout or all across that tip, or, occurred only along the back side of that tip”.

It is difficult to reconcile the prevailing model of budding in *Hornera* (Borg, 1926) with Boardman’s (1998) observations of two distinct types of autozooids, especially in species with wider exozones, such as *Hornera robusta*. Specifically, how can budding from the same basal budding locus form two distinct and *unbroken* layers (frontals and laterals) of autozooids? Drexler’s (1976) observations of widely distributed zooidal budding at the growing tip make more sense in this regard, but are at odds with Borg’s model. To improve understanding of hornerid branch construction we here conduct an investigation of internal anatomy using a variety of imaging methods.

## 2. Materials and Methods

Micro-CT, SEM and light microscopy were used to examine the skeletal and soft tissues of nine cancellate cyclostome species: eight hornerids and one crisinid. A range of hornerid ancestrulae and small colonies (<4 mm), as well as larger, usually reproductively mature, hornerid colonies, were imaged. Table 1 summarises the taxa, sampling localities and methods used. New Zealand specimens were collected by dredge or epibenthic sled, principally by the research vessels R.V. *Polaris* II (University of Otago) and R.V *Tangaroa* (National Institute of Water and Atmospheric Research). Culturing of live colonies of Otago shelf, Stewart Island and Snares shelf specimens was undertaken at Portobello Marine Laboratory, University of Otago, Dunedin.

**Table 1:**
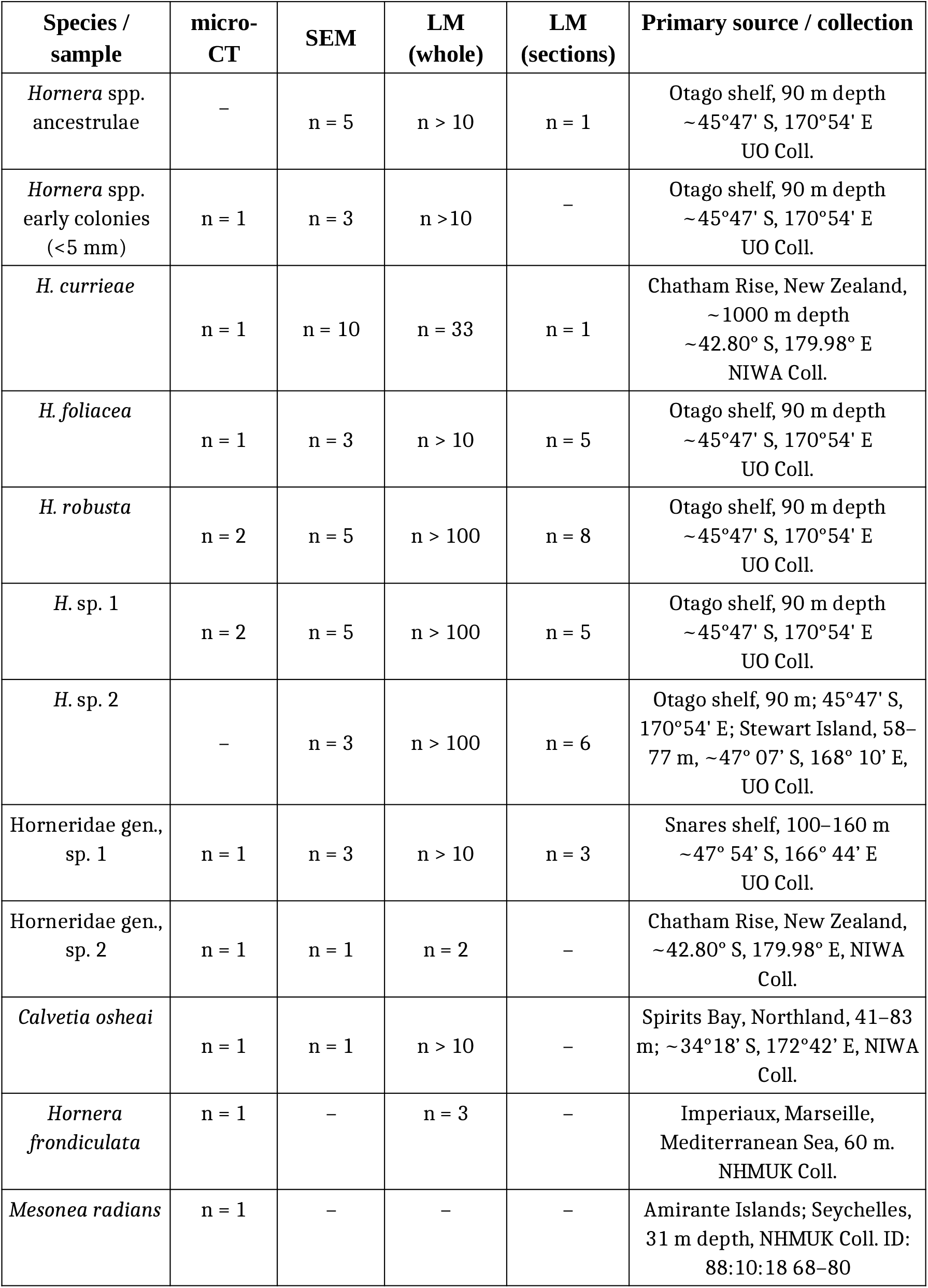
Material examined in this study. For micro-CT, SEM and light microscopy (LM) of live material, *n* gives the number of colonies/fragments imaged. In the case of sections, *n* is the number of examined semi-thin section sets. Institutional abbreviations: NHMUK, Natural History Museum, London, UK; NIWA, National Institute of Water and Atmospheric Research, New Zealand; UO, University of Otago, New Zealand.

We examined living lab-settled ancestrulae and small colonies, all identified herein as ‘*Hornera* sp.’ as it is currently difficult to identify these early stages to the species level (see Batson et al., 2019). Much larger colonies of *Hornera foliacea, Hornera robusta, Hornera* sp. 1 and *Hornera* sp. 2 were also maintained alive and imaged in culture. Collected colonies were transported to the laboratory and kept in a flow-through tank system in an isothermic room at ~ 13°C. Colonies were constantly supplied with natural particles and/or cultured algae via drip feeding or daily water changes. Methods used for laboratory culturing are described in more detail in Batson et al. (2019).

Regarding taxonomic identification, many of the New Zealand hornerids examined in the study are currently undescribed. All investigated morphotypes are well constrained, however, with species status confirmed by molecular sequencing by Andrea Waeschenbach and Helen Jenkins (NHMUK). A comprehensive taxonomic revision is underway (Abigail Smith et al., in preparation); readers are directed to that article for the eventual taxonomic identities of taxa studied in the present article. Voucher images of taxa studied herein are available by request.

### 2.1. Micro-CT (X-ray micro-computed tomography)

The primary subject of the autozooidal budding investigation was a small (~3 mm) colony of *Hornera* sp. growing on a colony of the cheilostome bryozoan *Hippomenella vellicata* from 90 m depth off Otago Peninsula (southeastern New Zealand, 45°47.890’ S, 170°54.50’ E). The basal part of this specimen was micro-CT scanned at high-resolution (~2 μm voxel size). Micro-CT scanning of mature colonies belonging to nine hornerid taxa was also conducted (Table 1). These included the following New Zealand taxa: (1) *Hornera currieae* Batson et al., 2021; (2) *Hornera foliacea* (MacGillivray, 1868); (3) *Hornera robusta* MacGillivray, 1883; (4) *Hornera* sp. 1; (5) Horneridae gen., sp. 1; (6) Horneridae gen., sp. 2; and (7) *Calvetia osheai* Taylor & Gordon, 2003. Micro-CT data was also available for *Hornera frondiculata* (Lamarck, 1816) from the Mediterranean, and the crisinid cancellate *Mesonea radians* from the Seychelles.

Micro-CT on New Zealand specimens was conducted using a Skyscan 1172 high-resolution micro-CT scanner (Bruker-MicroCT, Kartuizersweg. 3B, 2550, Kontich, Belgium). Organic matter was first removed from colonies using bleach solution (31.5 g.l^−1^ sodium hypochlorite) for 12 or more hours, followed by freshwater rinses, and oven drying at 60° C. Specimens were scanned at various resolutions yielding effective voxel sizes of ~ 1.8–3.4 μm. Data from the Mediterranean and Seychelles specimens was obtained using a Gatan XuM nanotomography system installed on a FEI Quanta 650 ESEM FEG scanning electron microscope at the NHMUK.

During scanning of the New Zealand material the beam-hardening correction was set to 100% and the aluminium and/or copper filter (0.5 mm) was engaged to further reduce beam-hardening artifacts; ring correction was set to 10. Operating voltage was 49–60 kV and amperage set to 200 mA. Specimen rotation was 180°, with rotation steps of 0.3°. *NRecon* (v.1.6.10.2) software was used for slice reconstruction. All New Zealand- and NHMUK-scanned datasets were processed in *FIJI* (v.2.0.0, US National Institute of Health), typically processed by conversion to 8-bit format and median filtering (2–3 px), and saved as TIF image sequences. Resulting stacks were imported into *Aviso* (2019 release, ThermoScientific) on a high-performance computer (~500 Gb RAM; Geophysics Laboratory, Department of Geology, University of Otago).

After thresholding, data visualizations were generated using front-face and back-face isosurface rendering, sometimes adding volumetric fog as a visual proxy for carbonate (Aviso Volren module). Anaglyphic 3D rendering was employed to assist with visual interpretation. In some cases, to isolate zooidal tubes from smaller-volume cavities within the skeleton (e.g. cancelli) the Dilation/Erosion modules were used, combined with use of the Interactive Thresholding and Ambient Occlusion modules in Aviso. Image editing and integration of interior–exterior isosurface renders were done with *Adobe Photoshop CS6*.

### 2.2. Scanning election microscopy (SEM)

Branch tips of eight New Zealand hornerid species were examined using scanning electron microscopy (Table 1). Cuticle and soft tissues were first removed from colonies using bleach solution (31.5 g.l^−1^ sodium hypochlorite) for 12 or more hours, followed by freshwater rinses and drying. Samples were sputter coated with gold–palladium and scanned with a JEOL 6700F FE-SEM at the Microscale and Nanoscale Imaging unit (OMNI).

### 2.3. Light microscopy

Live specimens were imaged using an Olympus dissecting microscope equipped with a digital camera and *Q Image* software. Some material was also fixed, either at the time of collection or shortly afterwards, and processed for sectioning. An unidentified ancestrula as well as branches from large colonies of five species (Table 1) were fixed with 2.5% glutaraldehyde solution in 0.1 mol^−1^ PBS with added sucrose (final solutions 990–1100 mOsmol) and processed using standard TEM protocol. Material was rinsed in 0.1 mol L^−1^ PBS with sucrose (same total osmolarity as fixative), decalcified with 10% buffered EDTA for 9–48 h and transferred into 1% OsO4 for 1.5 h. After osmication, the material was washed, dehydrated through a graded ethanol series and pure acetone, and embedded in Embed 812 epoxy resin. Specimens were sectioned with a diamond knife on Leica EM UC7 ultramicrotome and stained with toluidine blue or methylene blue. Resulting series of semi-thin sections (1 μm thick) were imaged with a light microscope (several Zeiss and Olympus models). In addition, we consulted four serial block-face SEM datasets obtained previously for *H. robusta* (see Tamberg et al., 2021 for details).

## 3. Results

### 3.1. Morphogenetic sequence of autozooidal budding

The micro-CT datasets reveal that budding of frontal and lateral autozooids is entirely decoupled. These two zooid types arise via two different budding modes taking place in two spatially separated budding sites. All hornerid taxa examined conformed to this dual-budding pattern (nine species, three genera).

Origins of the respective budding modes of frontals and laterals can be traced back to early astogeny, and are more easily understood in this light. Accordingly, the description below begins with the ancestrula and traces development of the basal stem, branch crown and the formation of the first branches. It is based on the high-resolution micro-CT scan of a small Otago shelf *Hornera* sp. colony (Figures 2, 3). Yielding about 1,000 slices, this dataset enabled detailed tracking of how morphogenetic events in the skeleton unfold, culminating in the distinctive unilaminate branch configuration of hornerids.

**Figure 2.**
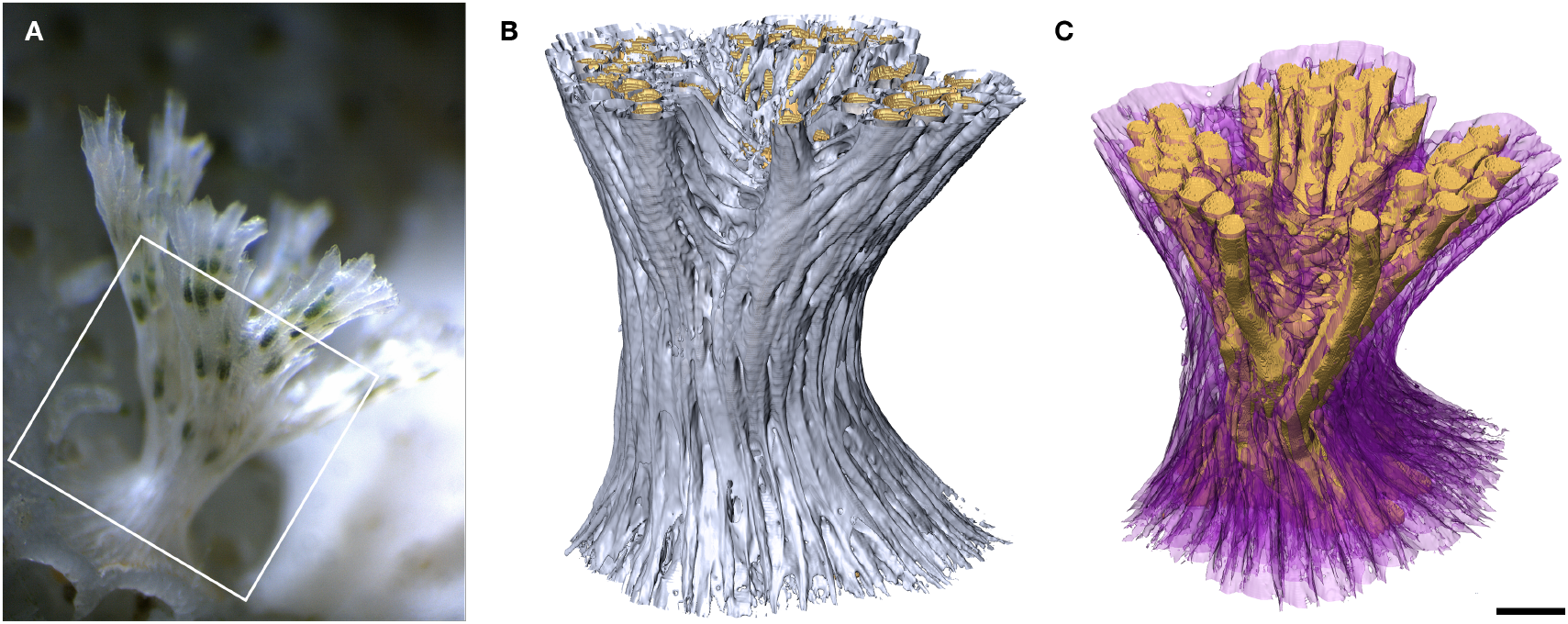
Early colony morphology of *Hornera* sp. **A.** Living colony of *Hornera* sp. of equivalent size to the micro-CT-scanned specimen used to study development of autozooidal budding. Box shows approximate region scanned. **B.** Exterior view of micro-CT-scanned basal stem and proximal part of the branch crown of *Hornera* sp. **C.** Interior view of same colony (back-face isosurface render). Autozooids digitally truncated; autozooids and large chambers formed by septa are coloured gold. Secondary calcification evident as a radial sequence of proximally directed kenozooids and cancelli (semi-transparent purple) originating from the autozooids. Scalebar ~200 μm.

**Figure 3.**
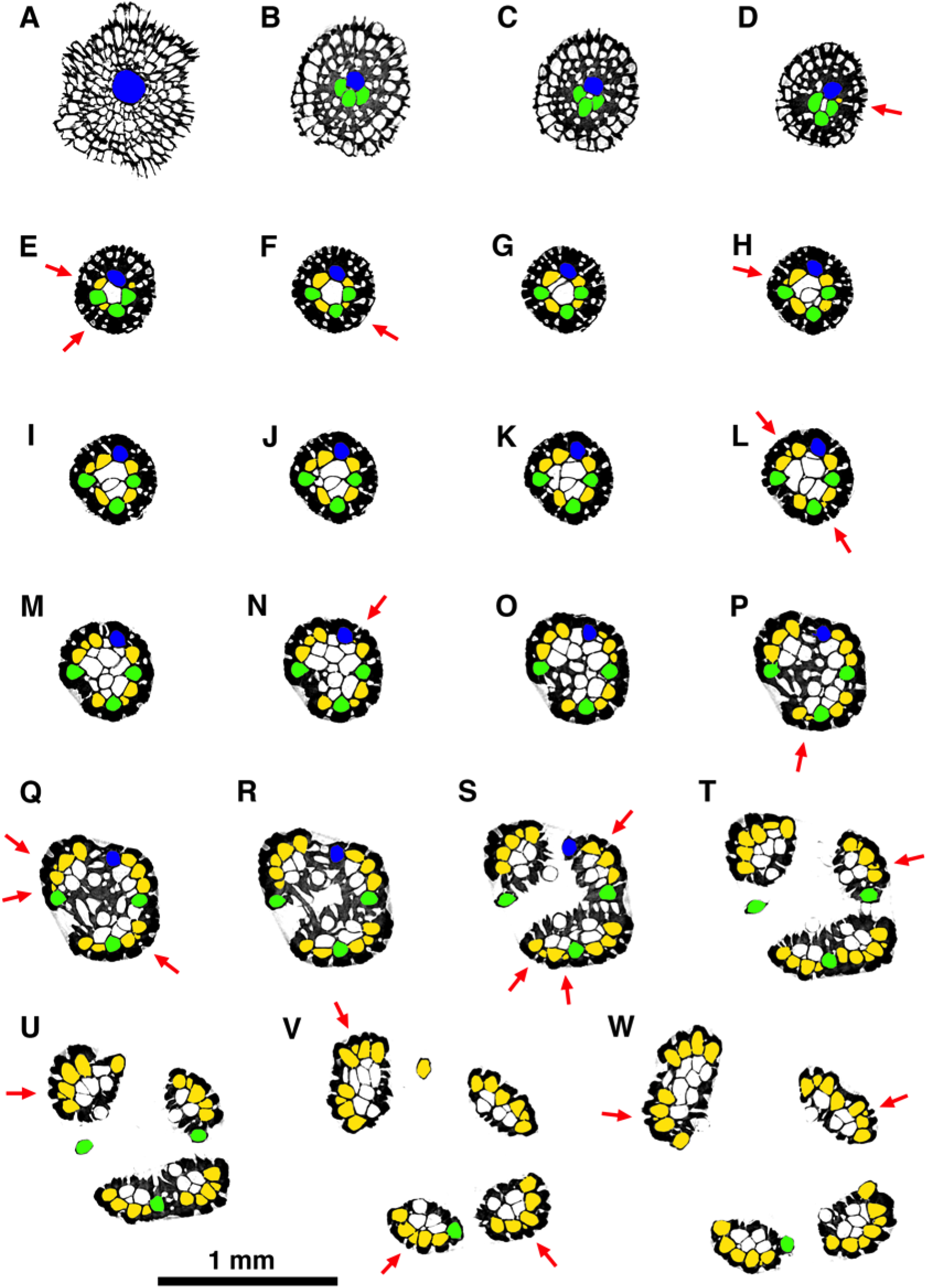
Sequence of selected micro-CT transverse orthoslices upwards through the ancestrula, basal stem and crown of a small colony of *Hornera* sp. Colours: blue, ancestrular zooid; green, periancestrular autozooids; yellow, exomurally budded lateral autozooids; white, frontal autozooids, kenozooids budded from the endozone and/or secondary kenozooids (outer layer). Each red arrow indicates the addition of a new exomurally budded lateral autozooid in the sequence. Sequence orthoslices A–W are discussed in main text.

The earliest autozooids are all laterals (Figures 3A–D, 4A–D). The first of these is, ultimately, the ancestrula itself (Figures 3A, 4A–D). The next laterals bud adventitiously onto the interior-walled roof of the ancestrula, forming a group of 1–7 periancestrular autozooids clustered around the ancestrular tube (Figures 3B–D, 4A–E). These early zooids typically (but not always—see Figure 4E) have fused skeletal walls and grow vertically, forming the basal stem. Periancestrular zooids are characterised by wide proximal footprints, about the same diameter as the autozooidal apertures of mature autozooids (Figure 4A–D), with the polypide buds located proximally (Figure 4B).

**Figure 4.**
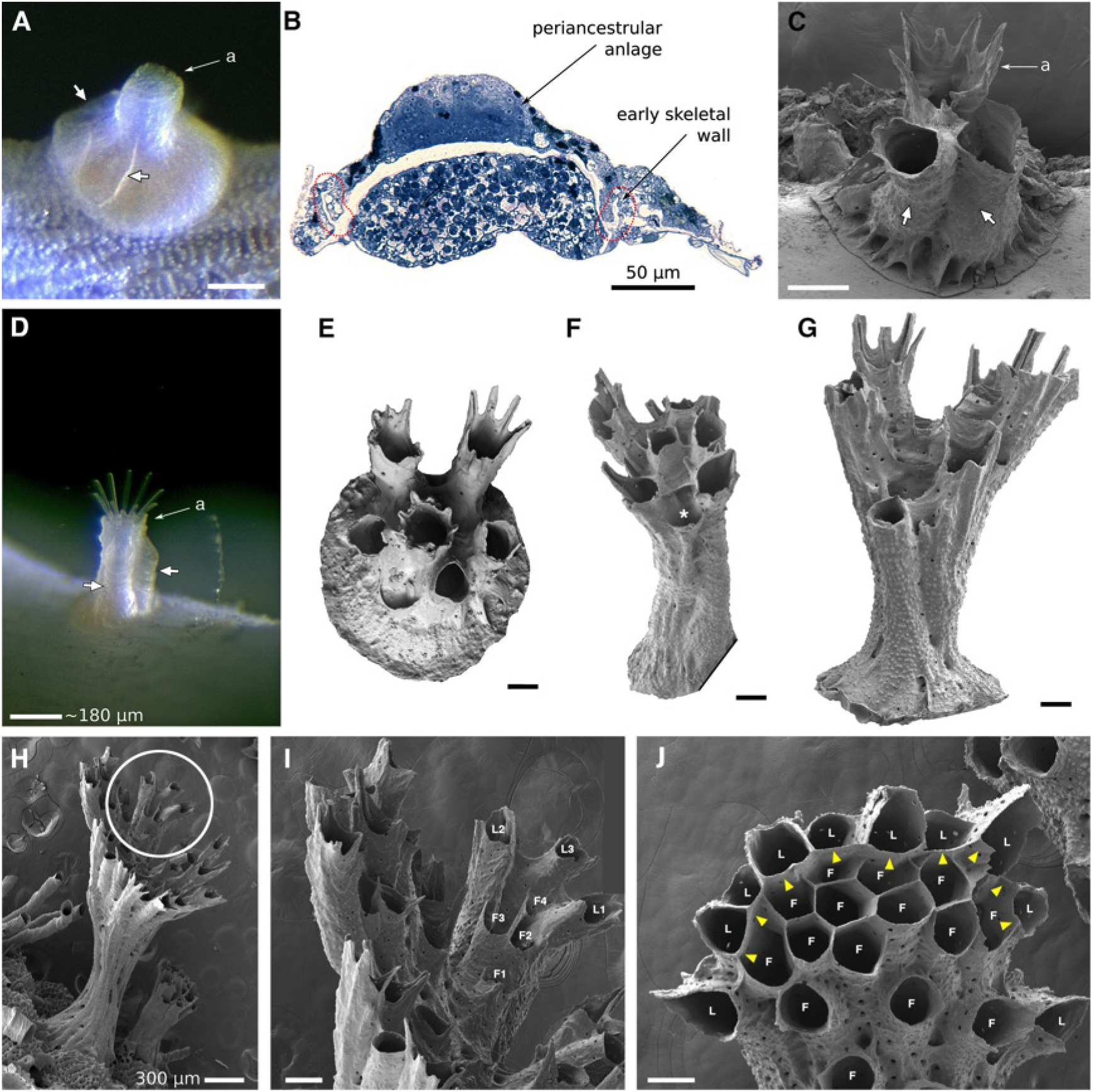
Astogeny in *Hornera* spp. **A.** Living ancestrula of *Hornera* sp., settled in the laboratory. White arrows show first-formed walls of two frontally budded adventitious (periancestrular) autozooids on ancestrula. Proximal footprint of each periancestrular zooid is approximately the same as a typical autozooid (a, aperture of ancestrula). **B.** Semithin section of the same ancestrula shown in A with the anlage of the periancestrular zooid atop the newly calcified ancestrular dome (longitudinal section, position roughly corresponding to short arrows in A). **C.** SEM of more-advanced ancestrula. White arrows show two periancestrular zooids partly fused with ancestrular tube (a). **D.** Live colony showing two periancestrular zooids (white arrows) growing up wall of central ancestrular zooid (a). **E.** Ancestrula of *?Hornera* sp. 2 from Foveaux Strait. Daughter autozooids are fully separated from each other. **F.** Multizooidal stem, with new exomurally budded lateral autozooid (asterisk). **G.** Incipient branch crown; diverging zooids are connected by septa, long zooids with peristomial spines are laterals; central space where frontal autozooids will bud centripetally is appearing. **H.** Young branch crown of *Hornera* sp. **I.** Close up of circled region in H, showing first-formed lateral autozooids (L1–L3); the frontal zooids (F1–F4) bud upon and grow along the frontal budding lamina formed by the basal surface of the laterals. **J.** Tip of mature branch of *Hornera* sp. Paramedial budding lamina is established (yellow arrowheads), frontals (F) and laterals (L) labelled. Note roofs of lateral autozooids alternate in size and distance from branch tip. Unlabelled scale bars, 100 μm.

As the stem extends vertically, additional lateral autozooids are budded coaxially onto the exterior of the distal stem by ‘exomural budding’ (new term: *exo*—outside, *mural*—wall). This type of zooidal budding takes place as discrete and spatially isolated budding events (section 3.2; Figures 3D–W, 4F–J, 5, 7A–F, 8). Autozooids and heterozooids can be budded onto the outer wall in this way.

**Figure 5.**
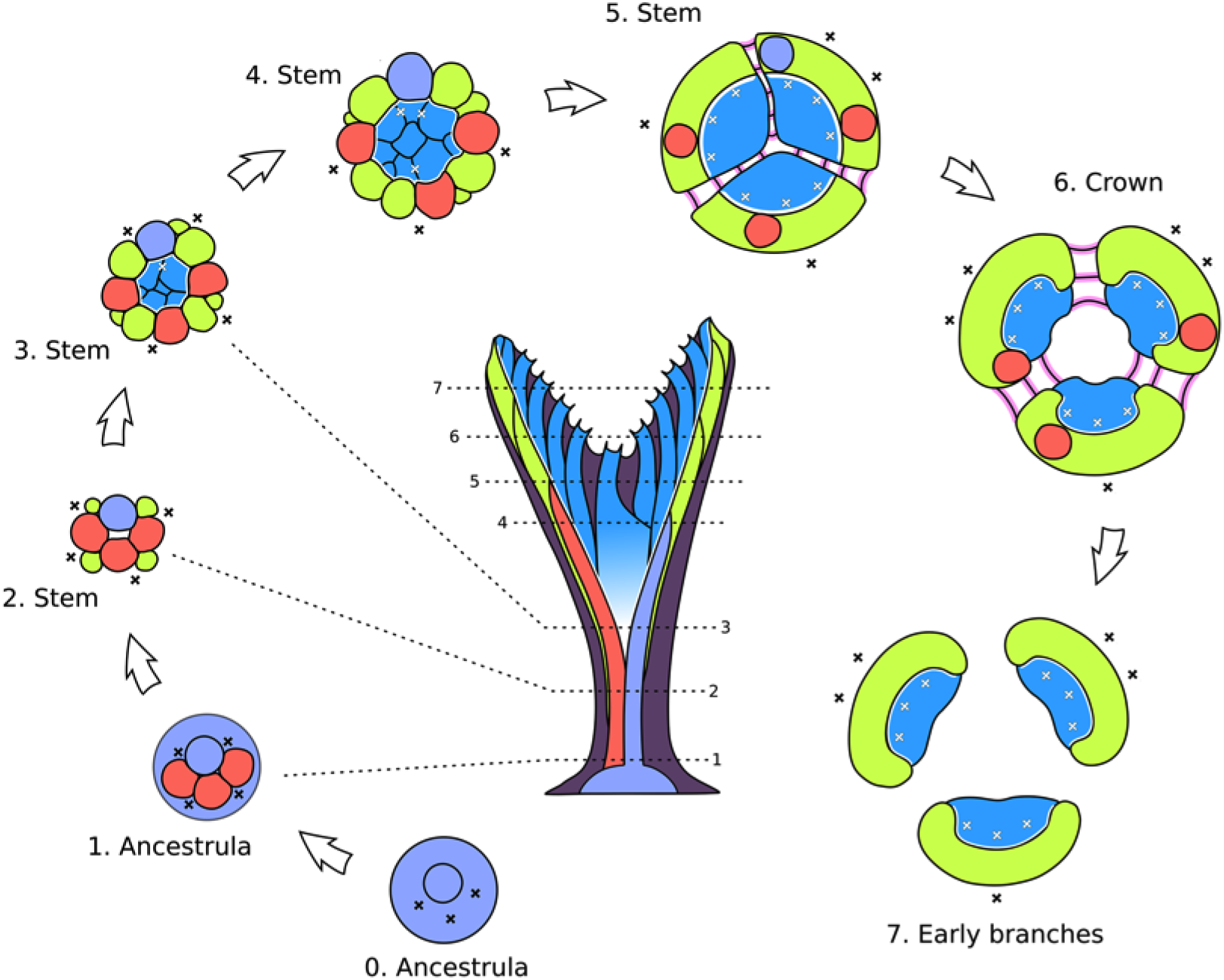
Formation of developmentally bilaminate branches from the ancestrula stage in *Hornera* sp. All numbered stages are transverse slices that show morphogenetic events occurring at or near the growing tip at the respective heights (dotted lines) of the colony shown in the central image—i.e., *not* cross-sections of a mature branch. **Lilac—** ancestrula; **red**—adventitious periancestular zooids; **green**—exomurally budded lateral autozooids; **blue**—basal budding of frontal autozooids from budding lamina; **pink**—septate budding of mostly kenozooids; **dark purple**—secondary wall calcification (shown in central image only). Black ‘X’, exomural budding locus; white ‘X’, budding events along budding lamina.

**Figure 6.**
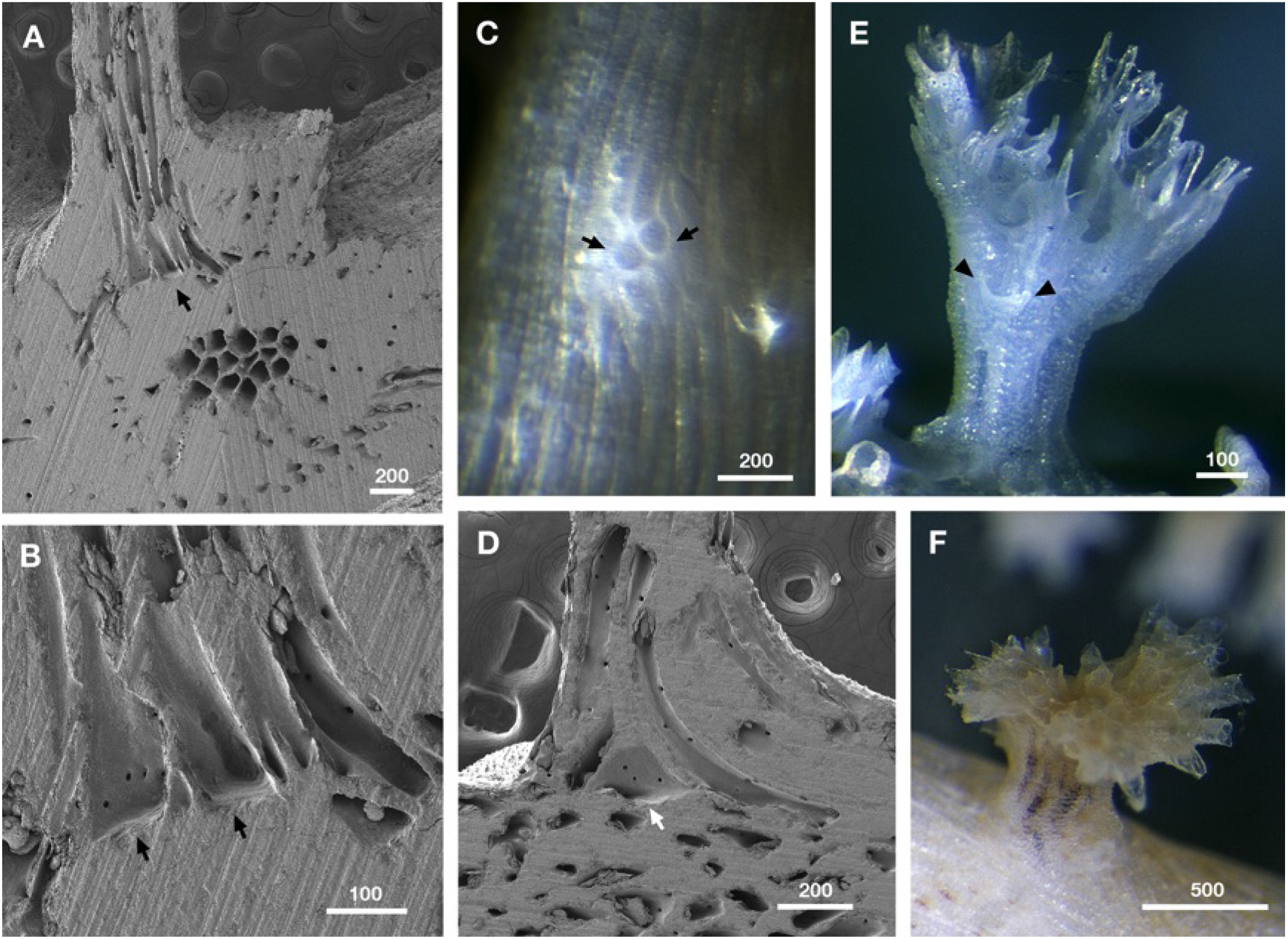
Secondary adventitious budding in *Hornera* spp. **A.** Longitudinally ground, adventitious, kenozooidal strut (top of image) budded on top of thick secondary wall of a transversely cut *Hornera* sp. 1 branch; abfrontal side up, strut growing perpendicular to branch surface (SEM). **B.** Close up of arrowed region in A. Two adventitiously budded kenozooidal chambers with flat bases corresponding to outer secondary skeletal wall arrowed. **C.** Adventitious zooid buds (arrowed) on abfrontal wall of living branch of *H. robusta*. Budding took place on a cultured branch resting directly on substrate, suggesting contact-induced budding of kenozooids. **D.** Large adventitiously budded chamber (arrow) at base of longitudinally ground kenozooidal strut, same orientation as A (*Hornera robusta*). **E.** Secondary adventitious or possibly delayed exomural budding on outer wall of basal stem (arrows). Focus-stacked micrograph of bleached colony. **F.** Adventitious branch crown with functioning autozooids (*H. robusta*). Scale bars in μm.

**Figure 7.**
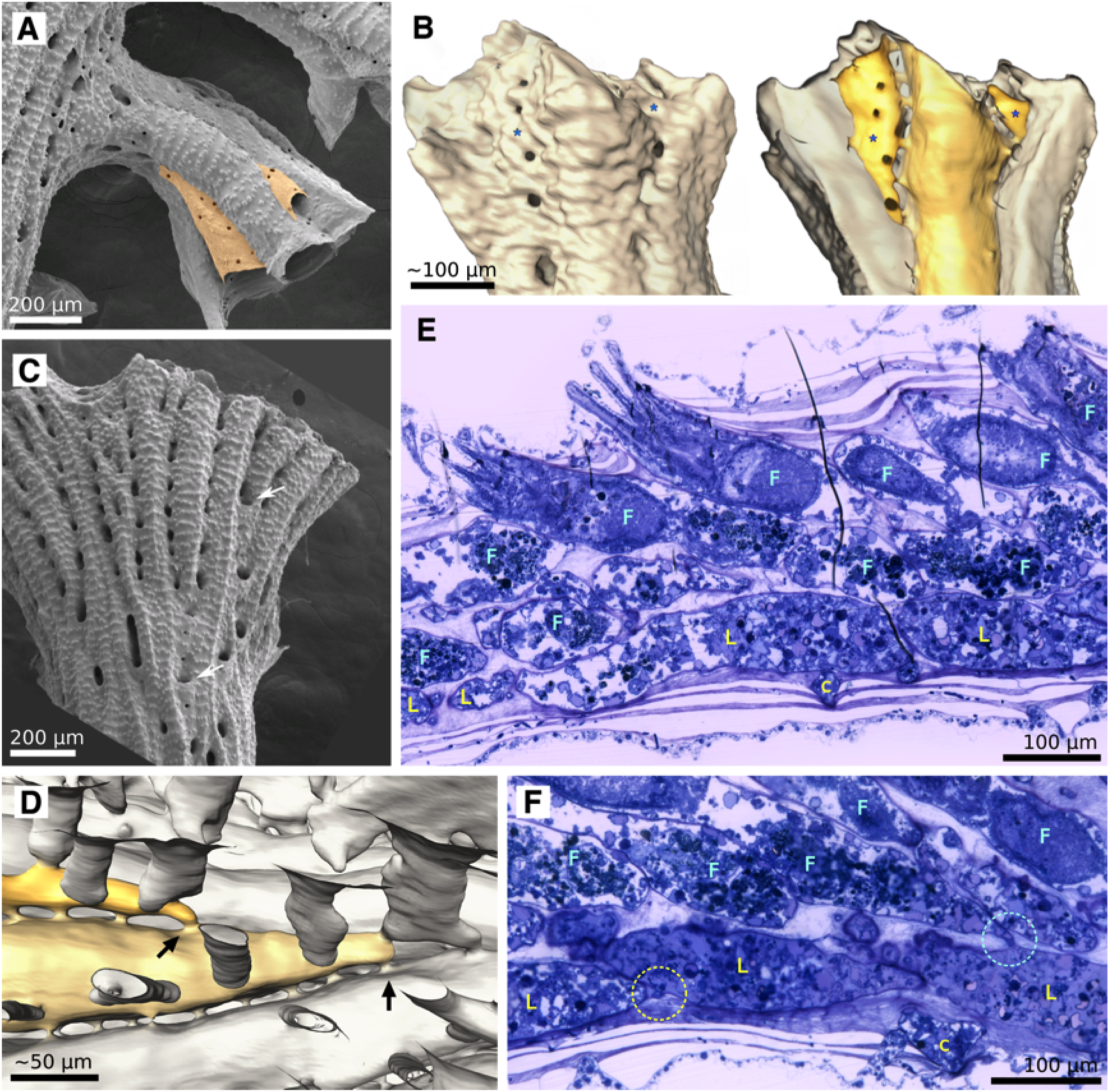
Exomural budding in *Hornera*. **A.** Conspicuous exomural budding of two autozooids (pale orange) on a secondarily formed (= lateral) branch of *Hornera robusta* (abfrontal view). **B.** Exterior (left) and interior (right) micro-CT reconstructions of the same *Hornera* sp. 1 branch tip showing two newly budded autozooids (*). Viewed externally, the location of zooids is obscured by secondary calcification. **C.** Abfrontal view of *H. robusta* branch tip, arrows show newly budded kenozooids in sulci, similar to, but more proximally sited, than exomurally budded autozooids. **D.** Hypostegal pore-associated exomural budding sites of two lateral autozooids of *Hornera* sp. 1; distal at left; some cancellus openings outlined for clarity. **E.** Sagittal semithin section of distal branch of *Hornera* sp. 2 showing bilaminate zooid arrangement. Lateral (L) and frontal (F) autozooids comprise two layers of separately budded chambers interfacing at paramedial budding lamina. **F.** Sagittal semithin section of *H*. sp. 2. Circle at left indicates hypostegal pore-associated exomural budding site of lateral autozooid; circle at right shows an interzooidal pore-associated budding site of frontal autozooid.

**Figure 8.**
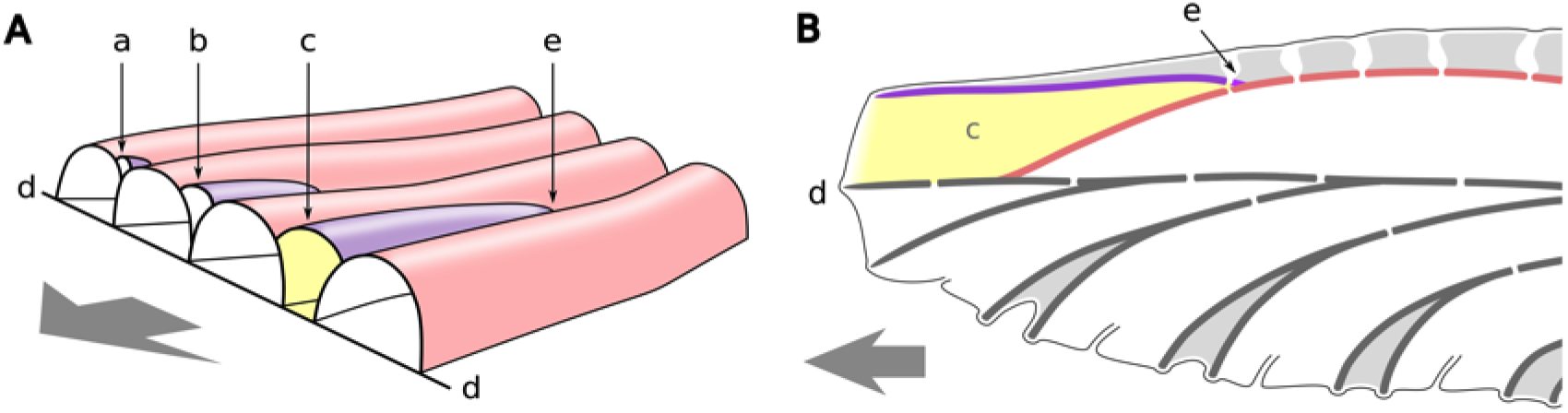
Schematic drawings showing two views of exomural budding and chamber development in the same *Hornera* branch (both drawings abfrontal side up). **A.** Changes in shape and position of newly budded lateral autozooid chambers (a–c) during distal growth and intercalation with older lateral autozooids (pink); (d) paramedial budding lamina; (e) exomural budding site. Frontal zooids not shown; arrow: distal. **B.** Longitudinal section of the yellow autozooid in A, with secondary calcification added. Colours and labels c, d, e correspond to A (additional colours: dark grey; primary calcification; light grey, secondary calcification).

Characteristically, each exomurally budded lateral autozooid bud: (1) is isolated and independent from other new lateral autozooid buds, (2) is centred upon (or in line with) a ‘hypostegal pore’ (see Batson et al., 2021) located in an outward-facing interzooidal sulcus (groove) (Figures 7–9), and (3) develops skeletal chamber walls at, or slightly proximally to, the growing tip—that is, a lag in wall calcification may be evident. Despite this, exomural budding normally takes place very close to the growing tip.

**Figure 9.**
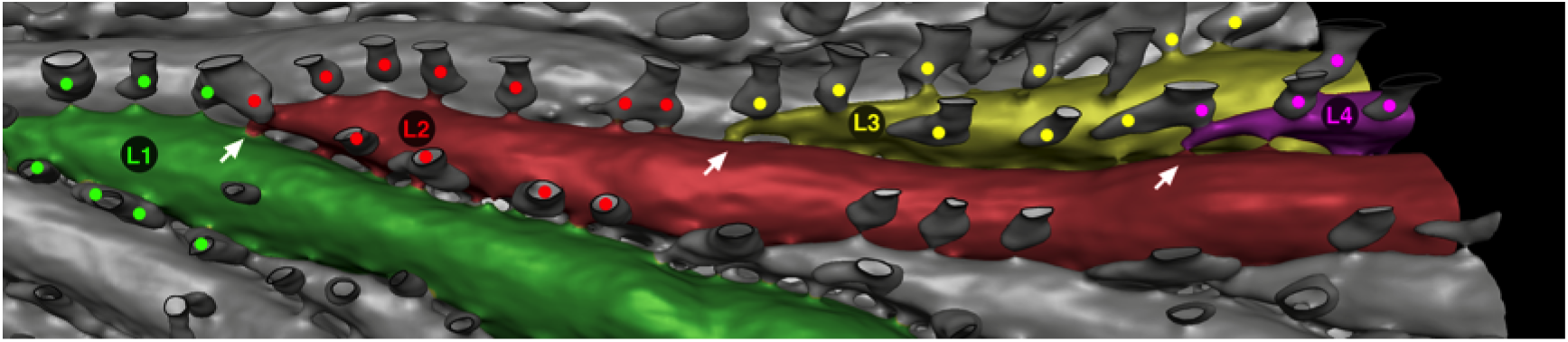
Sequence of lateral autozooid budding in mature branch of *Hornera* sp. 1 (abfrontal view, distal at right), showing four generations of autozooids (L1 to L4) and lines of associated cancelli (marked with dots). Cancelli are colour-coded to indicate from which autozooid its hypostegal pore originates within the sulcus (note: some cancelli connect to two zooids). Exomural budding sites marked by arrows. The cancelli arise from the most recently budded of the two zooids forming the sulcus, the transition taking place at the zooidal budding site. (Image: micro-CT back-face isosurface render; cancellus openings outlined with thin black lines to aid interpretation).

As they lengthen and widen, the newly budded lateral autozooids in the basal stem intercalate neatly with the existing zooids, forming an expanding ring of incipient laterals (Figures 3E–J, 5). Accompanying this process is the appearance and subsequent widening of the empty central region of the now-funnel-shaped stem. A circular budding lamina is soon established on the inner lining of this funnel, comprising the basal walls of the laterals. From this ‘ring budding lamina’ a series of roughly polygonal chambers is budded centripetally—the first of the future frontal autozooids (Figure 3G–P). This lamina remains unbroken until formation of the branch crown (Figures 3K–W, 4F–I).

Formation of the unilaminate branches occurs by splitting of the basal stem. In *Hornera* sp. this process is initiated around the time that the stem is ~ 1-mm-tall and consists of a core of ~ 10 future frontal autozooids surrounded by a ring of approximately the same number of laterals (Figures 3K, 4H). The stem splits radially into ~2–6 subequal divisions (three in the scanned specimen), which diverge from each other with continued growth to form the separate branches of the crown. Accompanying this process, septa form between the new branches, delineating the beginnings of chambers of future structural kenozooids in the axils (Figures 3K–Q, 4F, G).

During splitting of the basal stem, the ring budding lamina becomes fragmented and distributed among the daughter branches. Each section of the former ring becomes the paramedial budding lamina of the frontal autozooids in its own branch (Figure 3P–W). Immediately after splitting, the lateral autozooids ‘migrate’ around the axis of the now-separated branch to form a shallow ‘cup’ of lateral zooids when viewed in cross-section, such that the paramedial budding lamina and its frontal autozooids are enclosed on their lateral and abfrontal sides (Figures 3V–W, 4J, 5). These changes complete the ontogenetic sequence, setting in place the developmental template for all future budding in the branches of the colony.

### 3.2. Frontal budding in hornerids

Adventitious budding and exomural budding are two types of frontal budding. In *Hornera*, adventitious budding of autozooids occurs on the frontal wall of the ancestrula (Batson et al., 2019), whereas exomural budding takes place routinely on frontal walls of the stem and branch zooids. Occurrence of two, apparently rare, frontal budding modes in hornerids suggests potential homology, despite differences in the morphology of the proximal regions of their respective zooidal chambers. Zooidal chambers that appear morphologically intermediate between the two budding modes develop occasionally on the stem (e.g. Figure 6E).

Adventitious zooidal budding resembling the periancestrular budding seen on the ancestrula roof may also take place later in colony development. In some hornerid taxa (e.g. *Hornera robusta* and *H*. sp. 1), secondary zooids, kenozooidal struts and ‘reiterated’ branch crowns (Figure 6A–F) are formed in this manner. In addition, adventitious autozooidal budding is probably the origin of secondary ‘lateral branches’ commonly observed in older colony regions (see Harmelin, 2020, figure 2F).

Exomural budding gives rise to new lateral autozooids. Sites of exomural budding coincide with hypostegal pores located in the sulci lying between lateral autozooids. These pores connect the exosaccal cavity of an autozooid to the colony-enveloping hypostegal cavity. Exomural budding in *Hornera* always takes place in association with hypostegal pores, though these pores may be incipient at the time of budding. Away from the sulci, hypostegal pores are rare or absent on the frontal walls of lateral autozooids, possibly restricting budding opportunities. Within each sulcus the hypostegal pores are typically concentrated on one side of the groove—normally on the wall of the youngest of the two adjacent autozooids. Here they are more or less regularly spaced and form a line (Figures 7D, 9).

In the absence of exomural budding, hypostegal pores normally develop an associated cancellus (‘pore chamber’ of Borg, 1926), a tubular pocket of non-calcification in the secondary skeleton (Figures 7B, 9). However, during exomural budding, this pocket, or at least the site where such a pocket *would* develop, normally becomes the autozooidal budding locus (Figure 9). The sequence of morphogenetic events is uncertain and may be variable. It is unclear if the pocket—or even the primary skeletal wall and associated hypostegal pore(s)—have fully developed before morphogenesis of the polypide anlage takes place.

Micro-CT and soft-tissue sections show that the most-proximal part of a lateral autozooidal chamber usually coincides *directly* with a mural pore, suggesting the first skeletal walls may have formed around the pore (Figure 7D, F). In thin sections of mature autozooids these first pores are seen to be occupied by a pore cell (Figure 7F, and see Tamberg et al., 2022); although potentially these are formed secondarily, as cyclostome communication pores may be open early in ontogeny (Nekliudova et al., 2021, p. 20). If a pore is not located directly at the proximal-most tip of a zooidal chamber (i.e., the budding locus), one is invariably present very close by and normally within 5 μm.

Although the earliest stage of exomural budding of lateral autozooids was not observed directly, micro-CT and SEM of older chambers provide some information about early budding. They suggest that initial skeletal development involves growth of a hood-like structure where a sulcus begins to deepen due to zooid divergence. We were unable to determine whether the polypide bud develops in advance of the cystid frontal walls; however, this is a possibility given that the periancestrular autozooidal anlagen appear before skeletal walls develop (Figure 4B).

Exomural budding is followed by distal extension of the new zooidal chamber roof from the initial skeletal hood. The rate of chamber widening depends on the angle of divergence of the two underlying zooids forming the sulcus. At first the basal wall of the new lateral autozooid comprises the frontal walls of these two underlying laterals (Figure 8A, B). Once the underlying laterals have diverged fully, the new chamber meets the abfrontal surface of the paramedial budding lamina, which becomes the basal wall of the new lateral zooid (Figure 8A, B).

Timing of distal extension of exomurally budded zooidal chambers is variable. Frontal walls of new lateral autozooids can be slow to develop, sometimes lagging sufficiently far behind the growing tip to overgrow pustulose, secondarily calcified frontal walls (Figure 7A). In other cases, the leading edges of the frontal and basal walls grow together at the growing tip (Figure 7C). This variability may relate to a zooid’s stage of development, as well as its position within a colony (i.e. branch ‘leaders’ vs. lateral pinnules). In *Hornera robusta* the basal walls of lateral autozooids usually grow slightly in advance of the frontal autozooids in the main branches, such that the paramedial budding lamina is visible as a distinct leading edge (Figure 4J). This is particularly evident immediately after crown splitting, when the distal edges of frontal autozooid chambers are often set well back from the growing tip—Figure 4H, I).

The nature of hornerid branch construction by dual autozooidal budding modes is somewhat cryptic (e.g., Figure 7B–C). Examination of branch tips—often the only part of the colonial wall not covered by secondary calcification—can be misleading because of the way in which exomurally budded lateral autozooids grow. In SEMs of growing tips, it often appears that the lateral zooids have budded from the opposite side of the same budding lamina that gives rise to the frontal autozooids (Figure 4J). However, the frontal walls of these lateral autozooids merely ‘arrive’ at the budding lamina later in their development, having been budded onto frontal body walls (roofs) of existing laterals well away from the lamina (Figure 8). Newly budded lateral zooids may be mistaken for kenozooids (cf. Figure 7C).

Owing to the unusual three-dimensional growth trajectories of frontal and lateral zooids, zooidal budding as revealed by sagittal soft-tissue and skeletal sections can be difficult to interpret. This explains the contrasting accounts of Canu & Bassler (1920, p. 796) and Borg (1926, p. 306) compared to this study. Often all autozooids appear directed towards the frontal surface of the branch (Fig. 7E), as would be expected from a single basal budding lamina (i.e. in accordance with Borg’s 1926 budding model). Relatively few soft-tissue sections show unequivocal budding from the paramedial budding lamina or exomural budding (but see Figure 7F).

The dual modes of budding are responsible for some of the observed dimorphism between the frontal and lateral autozooidal chambers. In some *Hornera* species the lateral autozooids are considerably larger in diameter than frontals (see next section), and differ in cross-sectional shape, initially having wedge-shaped chambers, followed by ovoid or D-shaped chambers later in development. In contrast, lamina-budded frontal autozooids tend to be transversely polygonal within the endozone of wide-branched species, transitioning to subcircular after separating from the zooidal bundle (Figure 10).

**Figure 10.**
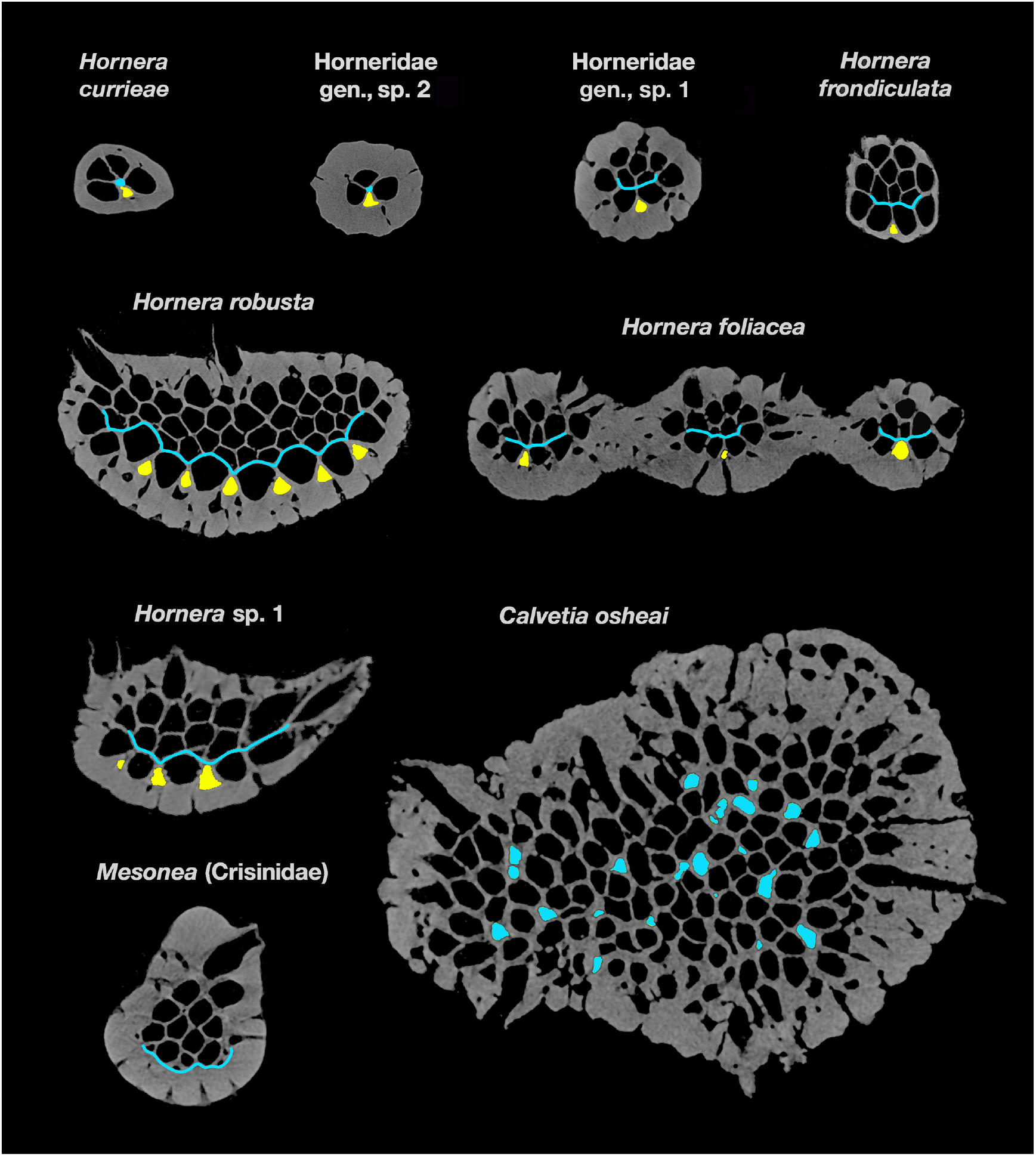
Micro-CT slices showing budding loci in main branches of eight hornerid species and one crisinid. Arrows show exomural budding loci of lateral autozooids; chambers of recently budded laterals are highlighted in yellow. Gracile taxa (*Hornera currieae*, Horneridae gen., sp. 1 & 2, and the fenestrate *H. foliacea*) each have a single medial budding locus for laterals; *Hornera robusta* and *H*. sp. 1 have multiple budding sites for laterals, and grow wider branches. Budding sites for frontal autozooids are shown in light blue. In *H. currieae* and Horneridae gen., sp. 1, budding site is axial; all other hornerids have paramedial budding laminae. In *Calvetia osheai* the main branches have widespread, unlocalised endozonal budding (possible newly budded chambers highlighted in blue). The crisinid *Mesonea* buds all autozooids from a basal budding lamina. Note: the thick secondary calcification was not present at the time of budding. Slices not shown to scale for clarity; slices edited to enhance contrast and remove non-skeletal material.

Lateral zooids in *Hornera* are typically longer on average than frontal zooids, and are more variable in length. In *H. robusta* and similarly wide-branched taxa, some laterals can be several millimetres long (Figure 1D). These persistent zooids tend to remain at or close to the abfrontal midline. They likely correspond to the ‘elongated tube inside the cortical part of the dorsal side’ reported to be the main budding locus in *H. antarctica* by Hennig (1911, p. 37). In other hornerid taxa, such as *H. foliacea* and *H. currieae*, the lateral autozooids bud and grow in a predictable herringbone (alternating) fashion, and zooidal chambers are typically more uniform in length.

### 3.3. Variability in autozooidal budding across taxa

In addition to nine hornerid cancellates, we examined the crisinid cancellate *Mesonea*. All the hornerid taxa had decoupled budding of frontal and lateral autozooids, although with some variations. In contrast, only a single zooidal budding mode was evident in *Mesonea*. Hornerid taxa below listed in order of cross-sectional size.

- *Hornera currieae*. This gracile hornerid has just two lines of frontal autozooids flanked by a line of laterals on each side (Figure 10). It has a small endozone with only 4–6 chambers in transverse slices. Consequently, a distinct budding lamina does not develop. However, the basic budding pattern is the same as in other hornerids—frontals (F) budded from the interior of the endozone at the interface with the laterals, and laterals (L) budded exomurally between diverging laterals.
- Horneridae gen., sp. 2. This undescribed New Zealand hornerid has the same 2F x 2L budding arrangement as *H. currieae*, and its budding proceeds similarly (Figure 10).
- Horneridae gen., sp. 1. The curved form of this gracile morphotype has 2–3 lines of frontal autozooids and a small frontal budding lamina (Figure 10). Exomural budding of laterals takes place solely from a median abfrontal budding locus.
- *Hornera frondiculata*. This European species has ~3–6 longitudinal rows of frontal autozooids (Harmelin, 2020). The scanned branch has three lines of autozooids and ~ 12–14 zooidal chambers visible in cross-sections; the well-developed paramedial budding lamina is in contact with up to four newly budded chambers in any given slice (Figure 10). Lateral autozooids are exomurally-budded along the abfrontal midline of the endozonal bundle in a strictly alternating pattern (one zooid curves left, next zooid curves right).
- *Hornera foliacea*. This fenestrate-branched hornerid has 2–4 rows of frontal autozooids (Figure 10). Autozooidal budding pattern is identical to previously covered taxa, despite the anastomosing growth. The micro-CT scan of this species covered a branch bifurcation, revealing how budding of lateral autozooids correlates with branching. Lateral autozooids bud along the branch midline; but, well before branching, the line of budding loci splits and diverges into two lines, the loci forming a Y-shaped arrangement when viewed abfrontally (Figure 11A). This split doubles the rate of lateral zooid budding, widening the branch well in advance of its bifurcation, whereupon each new branch has a single line of budding loci.
- *Hornera robusta*. This species has a very large endozone, especially in distal parts of large colonies, where branches can become wide and paddle-like. Transverse sections of main branches can contain more than 60 autozooidal chambers in various stages of development. MicroCT reveals that exomural budding is not confined to the branch midline, but occurs across most of the abfrontal branch surface (Figure 10). Not all of these incipient lateral chambers develop into lateral autozooids—at least some transition into kenozooids at branch bifurcations.

**Figure 11.**
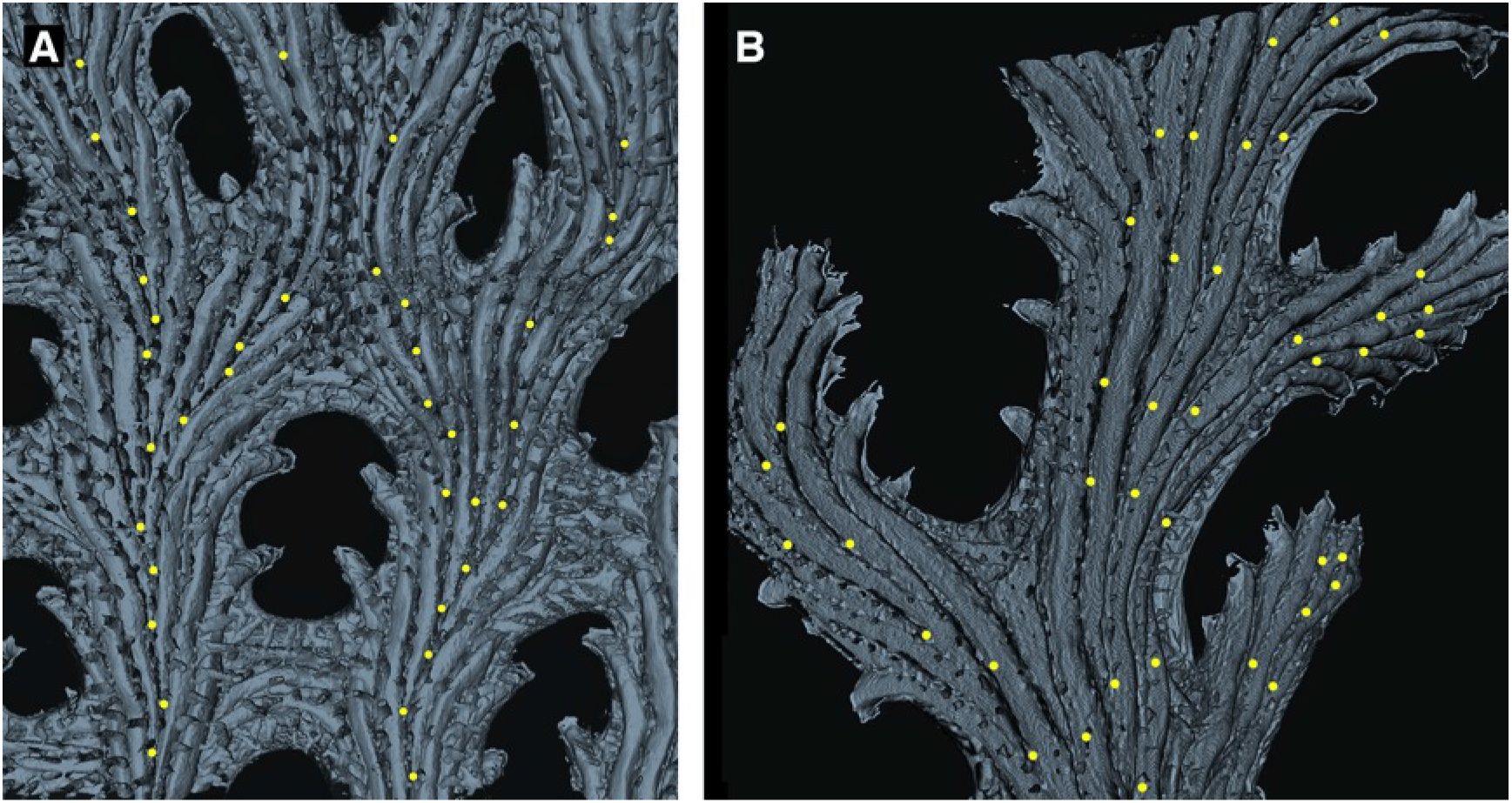
Interior skeletal reconstructions of exomural budding sites on abfrontal branch surface. Each budding locus is marked with a yellow dot. **A.** Part of fenestrate branch of *Hornera foliacea*. Note the difference in crossbar composition: the upper-central branch anastomosis results from branch fusion; the lower crossbar lacks exomural buds, having been formed solely by peristomial anastomosis. **B.** *Hornera*. sp. 1. Multiple lines of exomural budding sites associated with increased branch width. Denser concentrations of exomural buds result in shorter autozooids and higher zooidal divergence angles, evident in the pinnule on the centre-right. (Micro-CT-derived, back-face, isosurface renders).

Lateral multiplication of abfrontal budding sites is associated with branch widening in this and several other examined species (see Figure 11B), in contrast to taxa with lower cross-sectional zooid counts, such as *H. currieae* and *H. foliacea*, which are geometrically constrained to smaller branch size by their single lines of lateral autozooid budding sites. In terms of chamber dimorphism, *H. robusta* lateral zooids have significantly larger cross-sectional areas relative to frontal autozooids, a trait also seen in other hornerids (Figure 10).

- *Hornera* sp. 1. Exozones of this species are relatively thick, containing 20+ autozooidal chambers (Figure 10). The micro-CT-scanned branch has 3–4 parallel lines of lateral autozooid budding loci on the dominant branch and 1–3 lines on the side branches (Figure 11B).
- *Calvetia osheai*. Autozooids open on all sides of the thick cylindrical branches. Micro-CT shows seemingly chaotic clusters of unlocalised endozonal budding there (Figs. 10, 14). No consistent budding laminae were detected, although transient lamina-like features occur in places. Occasional eruptive unilaminate branches also develop (see Figure 14A–C), which show a hornerid-type budding pattern with distinct frontal budding lamina and abfrontal exomural budding of lateral autozooids (section 4.4). Lateral autozooids on unilaminate branches bud exomurally in multiple locations on the abfrontal surface, not only along the midline.
- *Mesonea* sp. Micro-CT of this non-hornerid cancellate reveals that all autozooids arise from a single basal budding lamina (Figure 10).

## 4. Discussion

The findings presented here differ from previous accounts of autozooidal budding in hornerids (Hennig, 1911; Canu & Bassler, 1920; Borg, 1926; 1944; Drexler, 1976). However, examination of figured thin sections of *Hornera antarctica* prepared by Hennig (1911, plate 5, figures 9–11) and Borg (1926, plate 8, figure 49) show evidence of composite branch construction by dual budding modes in this species too. The differences among published accounts probably reflect the cryptic nature of exomural budding, rather than its absence in hornerid taxa studied by previous researchers. In the present study the availability of internal 3D reconstructions based on micro-CT data was instrumental for recognising the dual budding types. In our opinion, composite branch construction by dual budding modes is widespread among extant Horneridae; one exception is an undescribed hornerid from southern New Zealand, which seems to have no laterals at all (Smith et al., in preparation).

For extinct hornerids we can be less certain about the mode of branch construction. Available reproductions of Drexler’s (1976) figures of Eocene species of *Hornera* were of insufficient quality to confirm exomural budding. A longitudinal thin section of one species investigated by Drexler (*Hornera reteramae*) is depicted in McKinney et al. (1993, figure 5.1). This section does not show exomural budding directly, but does show an arrangement of autozooids consistent with the presence of dual budding modes.

### 4.1. Definition of exomural budding (new term)

Despite uncertainty about how exactly exomural budding occurs, various traits set it apart from other forms of frontal budding. Exomural budding in *Hornera* takes place during primary branch morphogenesis, at or near the branch tip, and through the skeletal wall relative to the parent zooid. This form of budding is highly stereotyped, occurring in predictable locations (cf. lateral branching in tubuliporines). Autozooids budded this way are coaxially oriented from an early stage, growing along the branch axis, not perpendicular to the parent zooid(s). Additionally, developing lateral autozooids intercalate neatly with underlying laterals as they extend, maintaining a layer that is always one zooid thick (cf. cheilostome frontal budding, in which multiple layers of zooids accumulate).

No existing term in the bryozoological lexicon adequately describes this mode of budding. The term ‘adventitious’ is unsuitable because the budding loci are distal and zooids formed this way are tightly integrated into primary branch morphogenesis. Similarly, use of ‘frontal budding’ is not ideal: frontal budding in cheilostomes proceeds very differently (see Lidgard, 1989). It would be unhelpful to apply this umbrella term to the budding observed in hornerids. ‘Exomural budding’ is therefore proposed to define a subtype of stenolaemate frontal budding that takes place: (1) at discrete, unconnected loci, (2) beyond the outer perimeter of the existing zooidal endozone, and (3) near the branch growing tip, as part of routine primary growth.

### 4.2. Implications of exomural budding and composite branch construction

Adventitious budding and exomural budding are subtypes of frontal budding. That is, they take place on the frontal walls of autozooidal chambers, the region of body wall interfacing with environment on the “exposed or orifice-bearing side of the colony” (Ryland & Hayward, 1977). Functionally, these surfaces correspond to the ‘roofs’ of the zooids—the opposite side of the chamber to the basal wall, or zooid floor (Taylor et al., 2014). Figure 12 shows an interpretation of skeletal wall identities in *Hornera* based our study of colony development.

**Figure 12.**
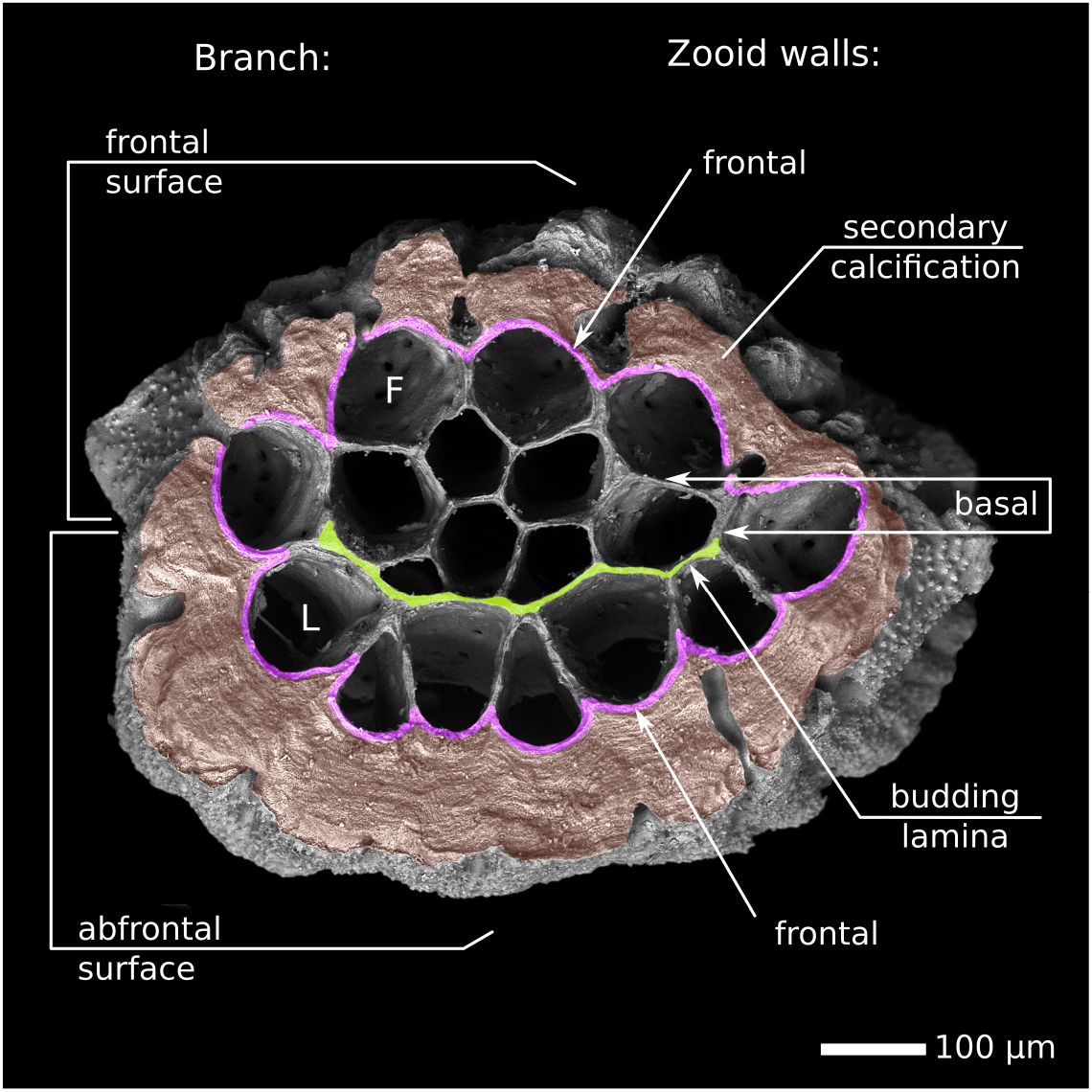
Respective orientations of branch and autozooidal walls in a branch segment of *Hornera* sp. 1 abscised by skeletal resorption (abfrontal side up). Frontal/lateral autozooidal chambers are dimorphic. Recently budded laterals are roughly triangular. The entire surface of the endozone is composed of developmentally frontal walls. SEM illuminated from below.

The two ‘outboard’ budding modes *in Hornera* are morphologically consequential for the colony. Firstly, they give rise to the peripheral ring of early lateral autozooids in the basal stem and all subsequent lateral autozooids in the colony. Secondly, the basal walls of the stem laterals coalesce to form the ‘ring budding lamina’, which is subdivided in the crown to become each branch’s paramedial budding laminae, the origin of the frontal autozooids. Finally, exomural budding loci also appear to be the origin of most of the structural kenozooids in the secondary body walls of the colony base and branch axils (Batson et al., unpublished data).

All autozooids in a *Hornera* colony have outward-facing frontal walls with respect to the orientation of the zooidal chambers. This differs from the cancellate *Mesonea* (Figure 10) and other unilaminate cyclostome taxa, which have developmentally basal walls (= zooid floors) facing the colony exterior on the abfrontal side of branches. In *Hornera*, the unusual autozooid configuration raises the question of whether its branches are truly unilaminate, as they are generally thought to be.

According to Hageman (2003): “Colonies that grow as a single layer of individuals are *unilaminate* […]. Colonies that grow erect in two back-to-back layers are *bilaminate*.” By this definition, hornerid branches are developmentally bilaminate. This trait is not immediately obvious because zooids budded on the abfrontal surface grow around the branch in order to open on the other side.

A more functionally oriented interpretation of *unilaminate* and *bilaminate* configurations is based on whether autozooidal apertures open on one or both sides of a branch or sheet (e.g. McKinney & Jackson, 1989, p. 56). This trait has important implications for feeding currents and colony form (e.g. McKinney, 1986; Suárez Andrés & Wyse Jackson, 2015). In this context *Hornera* could be considered to be developmentally bilaminate but functionally unilaminate.

Frontal and lateral polypides in all sections of *Hornera* that we examined were in the same anal–abanal orientation with respect to the branch frontal–abfrontal axis. If hornerids are developmentally bilaminate, this observation leads to the inference that the polypides of lateral autozooids have undergone a 180° rotation within the autozooidal chamber. Another alternative—i.e. that the abfrontal colony exterior is composed of the basal walls of lateral autozooids—seems unlikely. This is because: (1) the first exomurally budded zooids in a colony originate from the frontal walls of clearly frontally budded periancestrular zooids, and (2) budding takes place *across* the wall, rather than by septum formation within the zooidal chamber as occurs in cyclostome budding from basal walls.

Two aspects of exomural budding and subsequent zooid development facilitate composite branch construction in *Hornera*. First, the locations and timing of exomural budding events are well-matched to the rate required to replace lateral autozooids as they peel away from the endozone and open fronto-laterally. Secondly, distal growth of exomurally budded autozooidal chambers conforms closely to the divergence of the underlying autozooids. As the new lateral autozooid chambers lengthen, they also deepen, but their frontal walls (roofs) remain at the same level relative to the adjacent autozooids (Figures 8, 10). In this way, newly budded lateral autozooidal chambers gradually intercalate neatly into the series of surrounding autozooids, maintaining a single unbroken layer (Figures 4J, 5, 8). This highly orchestrated pattern of development differs from other known forms of frontal budding in cyclostome bryozoans.

How did dual autozooidal budding arise in hornerids? The answer may lie in the early development of the colony. In *Hornera*, adventitious budding of periancestrular autozooids establishes the frontally budded ‘lineage’ from which all lateral autozooids are descended (Figure 5). This suggests one scenario in which the periancestrular adventitious budding ability was co-opted and modified, becoming exomural budding. However, other evolutionary pathways are also possible, and it is presently unclear whether the ancestor of hornerids was bilaminate or unilaminate. Transitions from one branch configuration to another seem possible in bryozoans—for example, it has been argued that the unilaminate palaeostomate *Pseudohornera* evolved from a bilaminate ancestor (Tavener-Smith, 1975). At this point, it is best to wait until phylogenetic relationships, and the budding modes and early astogeny, both within and beyond Cancellata, have been better elucidated.

In this study we found a close association between hypostegal pores and the origins of lateral autozooids. It is unclear if pores have any direct role in the morphogenesis of the polypide anlagen—e.g., as a source of, or conduit for, blastemic cells. This may be unlikely given what is known about cyclostome budding, and its tendency to take place in the open physiological environment of the ‘common bud’ (Borg, 1926). However, the possibility is worth investigating considering that cheilostome pore chambers (mural pore analogues) are homologous with—and sometimes function as—autozooidal buds (Banta, 1969). Another possibility is that hornerid pores have a nutritional role for the developing (non-feeding) polypide (see Batson et al., 2021). For instance, interzooidal pores, in combination with mesothelial cell networks, have recently been shown to function as a *de facto* funiculus system in crisiids (Nekliudova et al., 2021).

The respective growth patterns of lateral and frontal autozooids may influence a well-known peculiarity of hornerid reproduction. In all *Hornera* species in which gonozooids have been studied, these female polymorphs are always derived from a frontal autozooid (e.g. Borg, 1944; Schäfer, 1991; Harmelin, 2020). This occurs despite the fact that lateral autozooids are volumetrically larger and located closer to the abfrontal branch surface, upon which the inflated incubation chamber develops. In *Hornera* this arrangement necessitates the growth of a tubular calcified conduit from the frontally located proximal part of the gonozooid, laterally around the outside of the branch and onto the abfrontal wall, where it expands into the incubation chamber.

The abfrontal position of the incubation chamber probably functions to avoid overgrowth of feeding zooids on the front of branches. But why do hornerids develop gonozooids and their associated abfrontal incubation chambers from frontal autozooids? Delayed development of reproduction relative to branch growth may be a factor. In all studied taxa, the gonozooids develop well behind the growing tip (e.g. Borg, 1926; Harmelin, 2020). Hornerids have progressive polypide cycling (Boardman, 1998), meaning that a polypide regenerates close to the zooidal aperture with each new degeneration–regeneration cycle. Via this iterated process, polypides of older lateral zooids migrate frontolaterally to the lateral sides of branches (Fig. 13), leaving the lateral autozooid chambers on the back of the branch mostly empty away from the tips, aside from brown bodies and sparse endocystal cell cover.

**Figure 13.**
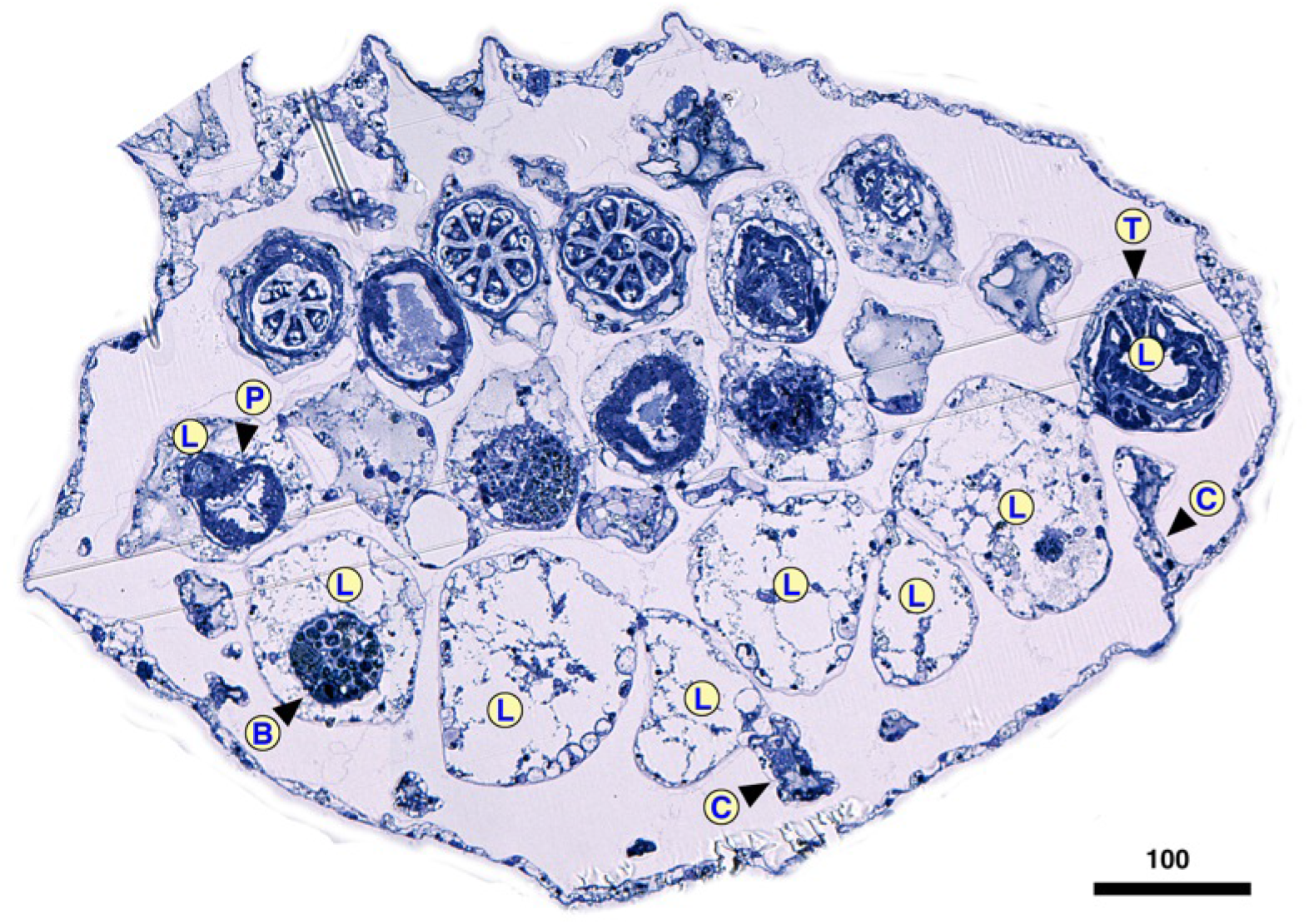
Transverse mid-branch section of *Hornera* cf. *robusta*, frontal side up. On the abfrontal side the lateral autozooid chambers (L) are empty, except for residual brown bodies (B) and sparse endocystal cell cover. Away from branch tips, polypides of lateral autozooids are located only at the lateral edges of the branch in wide-branched hornerid taxa. Note significant difference in frontal and lateral autozooid diameter. P, pharynx; T, tentacle; C, cancellus. Scale bar in μm.

Potentially, older lateral autozooids opening onto the sides of branches could develop into gonozooids, but this does not occur in any of the wide or narrow-branched taxa we examined. This observation raises several other possibilities: (1) lateral autozooids, whatever their position, are reproductively suboptimal for some other reason; or (2) hornerid ancestors may have lacked exomural budding and lateral autozooids, and the development of gonozooids from frontal autozooids could be a legacy of this phase of their evolution.

### 4.3. Comparison with other cyclostomes

How does autozooidal budding in hornerids compare with other cyclostome clades? Autozooids in the Cyclostomatida usually arise from the basal walls of the parent zooid or from a multizooidal lamina composed of incipient basal walls (Table 2). Articulates, tubuliporines, rectangulates, cerioporines and cinctiporids all bud autozooids in this fashion, as do cancellates—including the frontal autozooids of hornerids. Much less common is budding from frontal body walls. In cyclostomes the two known types of ‘frontal budding’ are adventitious budding and exomural budding. In both cases, the budding locus is the frontal wall of one or more parent zooids, though each type takes place in very different circumstances.

**Table 2.** Autozooidal budding types and subtypes reported across the Cyclostomatida (note, some overlap between categories). * Inferred from adnate phase of development.

| Autozooidal budding type | Skeletal wall type | Example taxa | Example colony region | Locus of budding on parent zooid | References |
| --- | --- | --- | --- | --- | --- |
| Intrazoooidal septum formation | Fixed walled taxa | Crisiidae<br>Oncousoeciidae (e.g. *Crisia*, *Crisulipora*, *Anguisia*) | Internodes; 'peristomial budding' | Basal * | Silén (1977); Taylor & Waeschenbach (2019); Ostrovsky (1998) |
| Budding from a multizoooidal basal lamina | Fixed and free walled | Tubuliporina<br>Cerioporina<br>Cancellata<br>Rectangulata (e.g. *Patinella*, *Hornera*) | Whole colony in many taxa or adnate stage in some erect taxa | Basal | Borg (1926); Hayward & Ryland (1985); Boardman (1998); This paper |
| Budding from within endozone | Fixed walled | Tubuliporina<br>Cerioporina (e.g. *Diaperoecia*, *Entalophora*) | Erect branches | Basal * | Borg (1926); Taylor et al. (2015) |
|  | Free walled | Cerioporines<br>Rectangulates (e.g. *Heteropora*) | Erect branches | Basal * | Taylor et al. (2015) |
| Intrazooecial 'fission' (budding) | Fixed walled | Tubuliporina (e.g. Eleidae) | Encrusting and erect branches | Vertical wall, | Taylor (1994) |
|  | Free walled | *Reptomulticava* (incertae sedis) | Encrusting | Vertical wall, possibly also frontal (?) | Hillmer et al. (1975); Nye & Lamone (1978) |
| Composite: endozonal & lamina budding in combination | Both wall types in single colony | *Terebellaria ramosissima* (Tubuliporina) | Erect branches | Basal | Taylor (1978) |
| Centripetal budding from a circumferential ring lamina | Fixed walled | *Tubulipora pencillata* (Tubuliporina) | Pedunculate stalk | Basal | Hayward & Ryland (1985) |
|  | Free-walled | Hornerid frontal autozooids (e.g. *Hornera*) | Basal stem | Basal, reverse side of lateral autozooids | This paper |
| Adventitious budding<br>= lateral branching *sensu* Harmelin, 1976) | Fixed walled | Tubuliporines (e.g. *Annectocyma*, *Voigttopora*) | Lateral branching 'runner' or 'ribbon' colonies | Frontal | Harmelin (1976) |
|  | Free walled | Hornerids<br>Periancestrular budding (e.g. *Hornera*) | Ancestrula | Frontal | This paper<br>Batson et al. (2019) |
| Exomural budding onto branch exterior | Free walled | Hornerid lateral autozooids (e.g. *Hornera*) | Branches and basal stem | Frontal | This paper |

Adventitious budding takes place in encrusting tubuliporines, such as *Annectocyma, Harmelinopora* and *Voigtopora* (Table 2). It occurs sporadically and enables secondary ‘lateral branching’, supplementing the distal budding that normally takes place in encrusting ‘runner’ and ‘ribbon’ colonies (Harmelin, 1976; Taylor et al., 2018). This budding mode is associated primarily with fixed-walled (exterior-walled) cyclostomes that lack a colony-enveloping epithelium; for this reason, skeletal resorption centred on pseudopores has been invoked as the likely mechanism by which a bud can emerge from the imperforate frontal wall (Harmelin, 1976). (Note: intrazooecial budding—‘fission’—*sensu* Hillmer et al. (1975) has also been regarded as a type of frontal budding as it results in overgrowth of the colony surface (e.g., Jablonski et al., 1997), but the budding locus is normally the vertical wall of the parent zooids, rather than the frontal wall *per se*.)

Within the suborder Cancellata, exomural budding is as yet only confirmed in the Horneridae. It would be illuminating to use micro-CT to examine the internal skeletal structure of other living cancellates—such as the pseudidmoneid *Pseudidmonea* Borg, 1944. This genus was recently shown to possess an interior-walled ancestrular roof and has a well-developed radiating multi-branched crown (Di Martino & Taylor, 2019) but it is not known whether *Pseudidmonea* has a zooidal budding pattern similar to hornerids. As for other currently valid cancellate families, Borg (1941) reported a different mode of early astogeny in the Crisinidae, and micro-CT confirms that *Mesonea* buds its autozooids solely from a basal budding lamina (Figure 10). The budding arrangement in Stigmatoechidae Brood, 1972 is unclear. It could be phylogenetically informative to examine thin sections of fossils of putative early hornerid genera: the Cretaceous genera *Eohornera* Brood, 1972 and *Siphodictyum* Lonsdale, 1849, as well as the Pliocene genus *Crassohornera* Waters, 1887.

Beyond Cancellata, the rectangulate family Alyonushkidae Grischenko et al., 2018 contains two genera of diminutive cyclostomes, *Alyonushka* and *Calyssopora*, that may perhaps undergo exomural budding of autozooids. It is noteworthy that *Alyonushka* possesses periancestrular zooidal budding—the only example of this type of budding known in a cyclostome outside Horneridae (Grischenko et al., 2018).

Frontal budding is uncommon among cyclostomes, and often appears to be used sparingly when present. In cheilostome bryozoans this type of budding is a phylogenetically widespread trait. An increase through time in the proportion of cheilostomes that undergo frontal budding was found by Lidgard (1989), with >60 % of Pliocene assemblages containing frontally budding encrusting taxa. One reason for the rarity of frontal budding in cyclostomes may be that other mechanisms for achieving ‘facultative self-overgrowth’ are available, such as the multizooidal budding laminae that overgrow older zooids in many cerioporines and rectangulates, and some tubuliporines (e.g., *Terebellaria*—Taylor, 1978).

### 4.4. Autozooidal budding in a cylindrically branched hornerid

The findings reported here may aid understanding of the morphology and evolution of the northern New Zealand hornerid, *Calvetia osheai* Taylor & Gordon, 2003. Unlike all other hornerids, this species has robust, roughly cylindrical, branches, with groups of autozooidal apertures opening on all sides (Figure 14A–E). Budding principally consists of apparently unlocalised endozonal budding at the tips of the main branches (Figures 10, 14E). Micro-CT and SEM conducted during the current study suggests that branches of *C. osheai* may be secondarily cylindrical, having been derived from a unilaminate (or potentially bilaminate) exomurally budding ancestor, possibly a species of *Hornera*.

**Figure 14.**
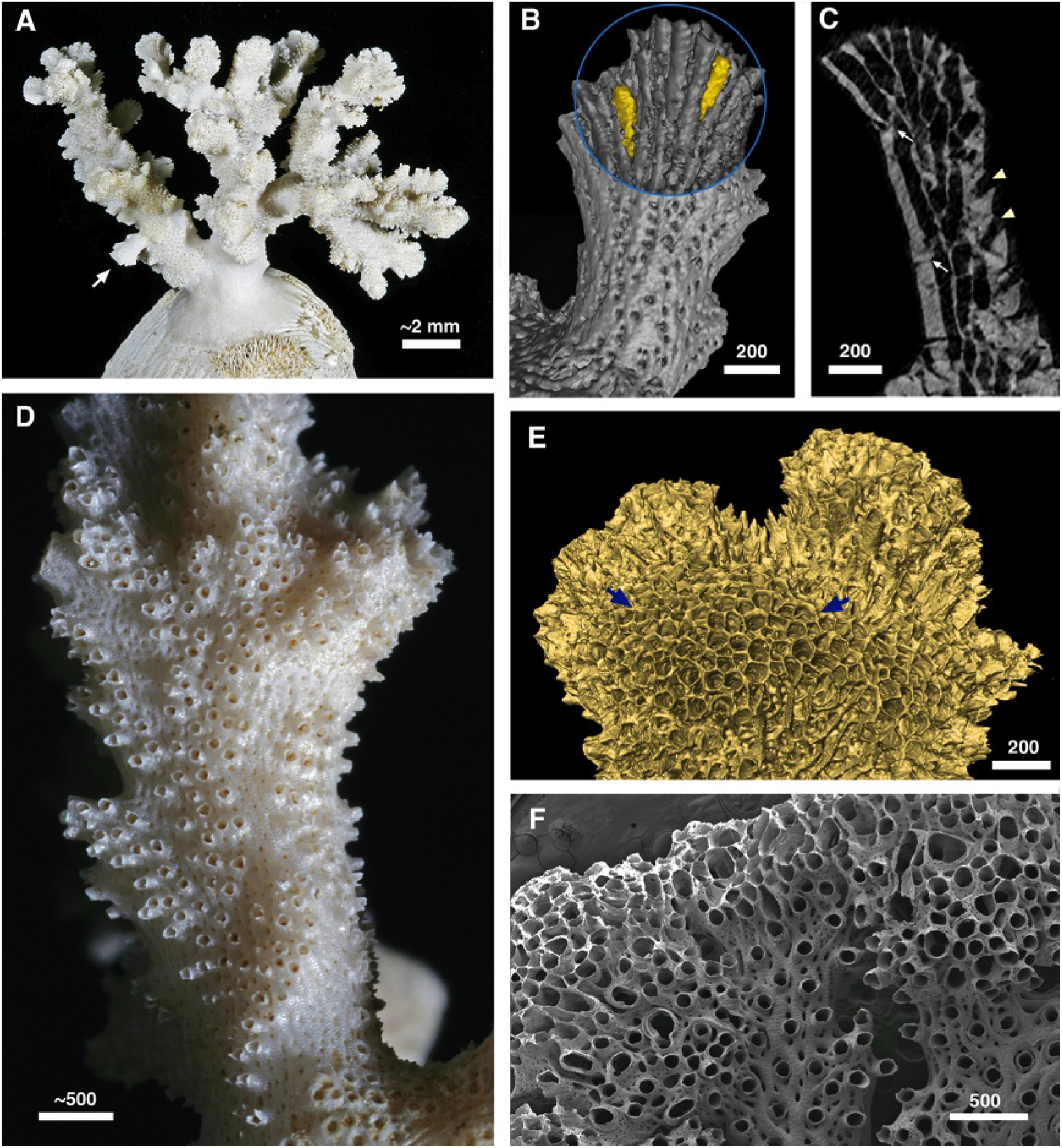
The hornerid *Calvetia osheai*. **A.** Colony growing on a bivalve shell; arrow indicates eruptive unilaminate branch. **B.** Combined interior/exterior isosurface render of the unilaminate branch arrowed in A, abfrontal view. Two exomurally budded ‘laterals’ highlighted in yellow. **C.** Longitudinal micro-CT slice of the same branch showing interior morphology: white arrows: exomurally budded lateral autozooids; arrowheads, apertures of ‘frontal’ autozooids. **D.** Branching pattern of autozooidal apertures, bordered by patches of zooid-free, cancellus-bearing wall, superimposed upon a much wider, roughly cylindrical branch of *C. osheai*. **E.** Branch tip, exterior view, with broad region of unlocalised budding, probably containing a mixture of autozooids and kenozooids (arrowed region). **F.** *Hornera* sp. 1. with teratological budding at branch tips, associated with local suppression of secondary calcification. Normal branch development visible at centre and bottom right. Scale bars in μm.

Evidence supporting a hornerid origin of *C. osheai* includes: (1) eruption of occasional short functionally unilaminate branches, with distinct frontal and lateral autozooids (Figure 14A–C); (2) lateral autozooids formed by exomural budding (Figure 14B, C); (3) localised development of narrow, haphazard, branch-like arrangements of autozooids seemingly ‘embedded’ within the broader, roughly cylindrical main branch surfaces, surrounded by cancellate walls (Figure 14D); and (4) in one case, an eruptive branch has a transversely oriented, incomplete skeletal abscission zone at its base (resorption-mediated branch abscission is a near-ubiquitous trait in other hornerids—Batson et al., 2020). If our interpretation is correct, the cancellus-bearing orifice-free walls surrounding the patches of autozooidal apertures may be homologous with the abfrontal wall of unilaminate hornerids.

We speculate that various developmental aberrations affecting zooidal budding in a *Hornera*-like ancestor, perhaps involving budding zone inversions, rotations and/or replications, could lead to the unusual morphology of *Calvetia*. Teratologies are sometimes present in *Hornera* in which secondary calcification becomes suppressed. When this occurs, disorganised kenozooids may develop between autozooids (Figure 14F), and the affected branches cease to grow much further. In *Calvetia*, the very ‘forgiving’ (i.e. self-organising and situationally plastic) secondary calcification typical of hornerids may facilitate long-term viability of the colonies by accommodating whatever peculiar budding transpires. In *C. osheai*, the patches of outer wall lacking autozooidal openings may function as exhalent channels for water currents generated in autozooidbearing areas.

*Calvetia* was originally assigned its own family, Calvetiidae, by Borg (1944), but its hornerid affinities were recognised by Taylor & Gordon (2003), who moved it to Horneridae. This may be an example of a genus-level transition in colonial growth form caused by a non-lethal disruption to zooidal budding. A second species of *Calvetia*, the Antarctic type species *C*. *dissimilis* Borg, 1944, bears a limited similarity to *C. osheai*. However, aspects of its surface structure resemble other Antarctic species—especially *Hornera smitti* Borg, 1944. This, combined with the unusual distributions of its species (one polar, the other subtropical—Taylor & Gordon, 2003), raises the possibility that *Calvetia* is diphyletic.

## Conclusions

Hornerid branches are here shown to have two distinct modes of autozooidal budding. These take place in two separate budding sites, and produce dimorphic zooidal chambers with different patterns of growth. Frontal autozooids bud from a paramedial budding lamina, whereas laterals bud from discrete sites onto the outer abfrontal branch wall by ‘exomural budding’, inferred to be a subtype of frontal budding. Exomural budding occurs in close association with hypostegal pores in the sulci of existing lateral autozooids at, or near, branch tips, and gives rise to coaxially oriented autozooidal chambers. Morphogenesis of these two zooid types occurs in parallel but is closely integrated. Together they form the unique composite structure that is the hornerid branch. Evidence from a range of taxa suggests that most extant hornerids employ dual budding modes during branch construction.

Early astogeny of a *Hornera* sp. colony is characterised by adventitious periancestrular budding onto the interior-walled ancestrula, which gives rise to the first lateral autozooids. Splitting of the basal stem at the branch crown leads to formation of the developmentally bilaminate, functionally unilaminate branches. Narrow-branched hornerid taxa possess a single line of medial budding sites on the abfrontal wall, whereas wide-branched species possess up to six, side-by-side, budding loci in locations on the abfrontal surface of branches.

Further study of the early stages of exomural budding is needed. The close association of autozooidal budding sites with mural pores in *Hornera* merits investigation. A wider survey of hornerid taxa, including putative Cretaceous and Cenozoic fossil taxa, would determine whether the dual autozooidal budding mode is a family-wide trait. If confirmed, this would provide new taxonomic characters for re-description of the family Horneridae, as well as hornerid genera and species. Cyclostome families suggested for future study of budding patterns include Stigmatoechidae, Pseudidmoneidae and the rectangulate family Alyonushkidae. Improved understanding of zooidal budding patterns in other cyclostomes may have utility as a source of new characters, especially in the light of new molecular phylogenies that do not agree with previous morphology-based classifications (Waeschenbach et al., 2009; 2012; Taylor et al., 2015).

## Acknowledgements

We thank the master and crew of R.V *Polaris* II, staff at Portobello Marine Laboratory and the Otago Microscale and Nanoscale Imaging unit (OMNI), University of Otago. We are grateful for technical support provided by Reuben Pooley, Linda Groenewegen, Andrew McNaughton, Hamish Bowman, Doug Mackie, Allan Mitchell, Sharon Lequeux and Liz Girvan (University of Otago). We also thank Mary Spencer Jones, Natural History Museum, London, UK and Jean-Georges Harmelin, Université Aix-Marseille, France. P.B. gratefully acknowledges Abigail Smith, Dennis Gordon and Daphne Lee for PhD supervision. P.B. received funding support from a University of Otago Doctoral Scholarship. Funding for computer infrastructure used to process and generate the micro-CT renders was provided by the Royal Society of New Zealand’s Rutherford Fellowship (16-UOO-001), the Ministry of Business and Innovation’s Endeavor Fund (C05X1605/GNS-MBIE00056), and the University of Otago’s ORG research grants program. In addition, we gratefully acknowledge access to a high-performance Titan X Pascal GPU kindly donated by the NVIDIA Corporation to the University of Otago Geophysics Laboratory.

## References

Banta, W. C. (1969). The body wall of cheilostome Bryozoa. II. Interzoidal communication organs. Journal of Morphology 129, 149–170.

Batson, P.B. & Probert, P. K. (2000). Bryozoan thickets off Otago Peninsula. New Zealand. Fisheries Assessment Report 2000/46. Ministry of Fisheries, Wellington, 31 pp.

Batson, P. B., Taylor, P. D. & Smith, A. M. (2019). Early astogeny in *Hornera* (Bryozoa; Cyclostomata; Cancellata). Australian Palaeontological Memoirs 52, pp. 23–30.

Batson, P. B., Tamberg, Y., Taylor, P. D., Gordon, D.P & Smith, A. M. (2020). Skeletal resorption in bryozoans: occurrence, function and recognition. Biological Reviews. 95, pp. 1341–1371. DOI: 10.1111/brv.12613.

Batson, P. B., Tamberg, Y., Gordon, D. P., Negrini, M., & Smith, A. M. (2021). *Hornera currieae* n. sp. (Cyclostomatida: Horneridae): a new bathyal cyclostome bryozoan with reproductively induced skeletal plasticity. Zootaxa 5020 (2): pp. 257–287. https://doi.org/10.11646/zootaxa.5020.2.2

Bock, P. E., & Gordon, D. P. (2013). Phylum Bryozoa Ehrenberg, 1831. Zootaxa, 3703(1), 67–74.

Boardman, R. S. (1998). Reflections on the morphology, anatomy, evolution, and classification of the class Stenolaemata (Bryozoa). Smithsonian Contributions to Paleobiology; no. 86, 59 p.

Boardman, R. S., & McKinney, F. K. (1985). Soft part characters in stenolaemate taxonomy. In C. Nielsen & G. P. Larwood (Eds.), Bryozoa: Ordovician to Recent (pp. 35–44). Olsen & Olsen.

Boardman, R.S, and Buttler, C. J. (2005). Zooids and extrazooidal skeleton in the order Trepostomata (Bryozoa). Journal of Paleontology, 79 (6): 1088–1104.

Borg, F. (1926). Studies on Recent cyclostomatous Bryozoa. Zoologicska Bidrag Från Uppsala. Band 10. Almqvist & Wiksells, Uppsala, 507 pp.

Borg. F. (1941). On the structure and relationships of Crisina (Bryozoa, Stenolaemata). Arkiv for Zoologi, 33A (11), 1–44.

Borg, F. (1944). The stenolaematous Bryozoa. In Bock, S., ed. Further Zoological Results of the Swedish Antarctic Expedition 1901-1903, 3, Stockholm: Norstedt and Söner, 1–276.

Borg, F. (1965). A comparative and phyletic study on fossil and Recent Bryozoa of the suborders Cyclostomata and Trepostomata. Arkiv for Zoologi, ser. 2, v. 17, no. 1, p. 1–91.

Canu, F. & Bassler, R. S. (1920). North American Early Tertiary Bryozoa. United States National Museum Bulletin 106:1–879, 279 figs., 162 pl.

Cheetham, A. H. (1986). Branching, biomechanics and bryozoan evolution. Proceedings of the Royal Society of London. Series B, Biological Sciences 228, 151–171.

Di Martino, E. & Taylor, P. D. (2019). *Pseudidmonea* Borg, 1944 (Cyclostomata: Pseudidmoneidae): description of two new species from the Miocene of New Zealand and phylogenetic relationships of the genus. Australasian Palaeontological Memoirs 52, 67–75. ISSN 2205–8877.

Drexler, W. W. (1976). Wilmington Jacksonian hornerids – important cyclostome bryozoans from the upper Eocene Castle Hayne Limestone in southeasternmost North Carolina. Unpublished dissertation. The Pennsylvania State University, College Park, 112 p.

Grischenko, A. V., Gordon, D. P. & Melnik, V. P. (2018). Bryozoa (Cyclostomata and Ctenostomata) from polymetallic nodules in the Russian exploration area, Clarion-Clipperton Zone, eastern Pacific Ocean—taxon novelty and implications of mining. Zootaxa, 4484 (1), 1–91. https://doi.org/10.11646/zootaxa.4484.1.1

Hageman, S. J. (2003). Complexity generated by iteration of hierarchical modules in Bryozoa. Integrative and Comparative Biology 43, 87–98.

Harmelin, J. G. (1976). Le sous-ordre des Tubuliporina (Bryozoaires Cyclostomes) en Méditerranée: écologie et systématique. Institut Océanographique, Fondation Albert Ier, Prince de Monaco.

Harmelin J.-G. (2020). The Mediterranean species of *Hornera* Lamouroux, 1821 (Bryozoa, Cyclostomata): reassessment of *H. frondiculata (Lamarck, 1816) and description of H. mediterranea n. sp*. Zoosystema 42 (27): 525–545. https://doi.org/10.5252/zoosystema2020v42a27

Hayward, P. J. & Ryland J. S. (1985). Cyclostome Bryozoans: Keys and notes for the identification of the species. E. J. Brill/Dr. W. Backhuys. London, 147 p., 48 fig.

Hennig, A. (1911). Le Conglomérat Pleistocène à Pecten de I’ile Cockburn. Wissenschaft. Ergebnisse der Schwed Südpolar Expedition 1901-1903, Stockholm, 3 Lief, pp. 1–72, pls. 5.

Hillmer, G. Gautier, T. G. & McKinney, F. K. (1975). Budding by intrazooecial fission in the stenolaemate bryozoans *Stenoporella*, *Reptomulticava* and Canalipora: Mitteilungen aus dem Geologisch-Palaontologisches Institut der Universitiit Hamburg, v. 44, p. 123–132.

Jablonski, D., Lidgard, S. & Taylor, P. D. (1997). Comparative ecology of bryozoan radiations: origin of novelties in cyclostomes and cheilostomes. Palaios 12, 505–523.

Lidgard, S. (1985). Budding process and geometry in encrusting cheilostome bryozoans. In: Nielsen C, Larwood GP (eds) Bryozoa: Ordovician to Recent. Olsen and Olsen, Fredensborg, pp 175–182.

Lidgard, S. & Jackson, J. B. C. (1989). Growth in encrusting cheilostome bryozoans: I. Evolutionary trends. Paleobiology 15: 255–282.

Ma, J.-Y., Buttler, C. J. and Taylor, P. D. 2014. Cladistic analysis of the ‘trepostome’ Suborder Esthonioporina and the systematics of Palaeozoic bryozoans. Studi Trentini di Scienze Naturali, 94, 153–161.

Mongereau, N. (1972). Le genre *Hornera* Lamouroux, en Europe (Bryozoa-Cyclostomata). Annalen des Naturhistorischen Museums, Wien, 76:311–373.

McKinney, F. K. (1986). Evolution of erect marine bryozoan faunas: repeated success of unilaminate species. American Naturalist 128, 795–809.

McKinney, F. K. & Jackson, J. B. C. (1989). Bryozoan Evolution. University of Chicago Press.

McKinney, F. K., Taylor, P. D., and Zullo, V. A. (1993). Lyre-shaped hornerid bryozoan colonies: homeomorphy in colony form between Paleozoic Fenestrata and Cenozoic Cyclostomata. Journal of Paleontology 67 (3): 343–354.

Nekliudova, U. A., Schwaha, T. F., Kotenko, O. N., Gruber, D., Cyran, N. & Ostrovsky, A. N. (2021). Three in one: evolution of viviparity, coenocytic placenta and polyembryony in cyclostome bryozoans. BMC Ecology & Evolution, 21, 54. https://doi.org/10.1186/s12862-021-01775-z

Nye, O. B. & Lamone, D. V. (1978). Multilamellar growth in *Reptomulticava texana*, a new species of cyclostome Bryozoa. Journal of Paleontology 52, 830–845.

Ostrovsky, A. N. (1998). The genus *Anguisia* as a model of a possible origin of erect growth in some Cyclostomatida (Bryozoa). Zoological Journal of the Linnaean Society, 124: 355–367.

Ryland, J. S. (1970). Bryozoans. Hutchinson University, London, 175 pp.

Ryland, J. S. and Hayward, P. J. (1977). British Anascan Bryozoans. Synopses of the British Fauna, 10, 1–188.

Schäfer, P. (1991). Brutkammern der Stenolaemata (Bryozoa): Konstructionsmorphologie und phylogenetische bedeutung. Courier Forschungsinstitut Senckenberg, 136: 147 pp.

Schwaha, T. F., Ostrovsky, A. N. & Wanninger, A. (2020). Key novelties in the evolution of the aquatic colonial phylum Bryozoa: evidence from soft body morphology. Biological Reviews, 95 (3), 696–729. https://doi.org/10.1111/brv.12583

Silén L. (1977). Structure of adnate colony portions in Crisiidae (Bryozoa Cyclostomata). Acta Zoologica, Stockholm, 58, 227–244.

Smith, A. M., Taylor, P. D. & Spencer, H. G. (2008). Resolution of taxonomic issues in the Horneridae (Bryozoa: Cyclostomata). In: Wyse Jackson, P. N. & Spencer Jones, M. E. (Eds.), Annals of Bryozoology. Vol. 2. Aspects of the History of Research on Bryozoans, International Bryozoology Association, Dublin, pp. 359–412.

Smith, A. M., Batson, P. B., Waeschenbach, A., Taylor, P. D., Jenkins, H. L., Gordon, D. P. (In Preparation). Molecular and morphological techniques show bryozoan family Horneridae (Cyclostomata: Cancellata) in southern New Zealand is unexpectedly diverse.

Smitt, F. A. (1867). Kritisk förteckning öfver Skandinaviens hafs-Bryozoer. Öfversigt af Kongliga Vetenskaps-Akademiens Förhandlingar 23: 395–534, pls 1–13.

Suárez Andrés, J. L. and Wyse Jackson, P. N. (2015). Feeding currents: a limiting factor for disparity of Palaeozoic fenestrate bryozoans. Palaeogeography, Palaeoclimatology, Palaeoecology 433, 219–232.

Tamberg, Y., Batson, P. B., & Napper, R. (2021). Polypide anatomy of hornerid bryozoans (Stenolaemata: Cyclostomatida). Journal of Morphology, 1–18. https://doi.org/10.1002/jmor.21415

Tamberg Y., Batson, P. B., Smith, A. M. (In press). The epithelial layers of the body wall in hornerid bryozoans (Stenolaemata: Cyclostomatida). Journal of Morphology (JMOR-21-0196).

Tavener-Smith, R. (1975). The phylogenetic affinities of fenestelloid bryozoans. Palaeontology 18:1–17, 5 fig., pl. 1–2.

Taylor, P. D. (1978). The spiral bryozoan *Terebellaria* from the Jurassic of southern England and Normandy, Palaeontology 21, 357–391.

Taylor, P. D. (2000). Cyclostome systematics: Phylogeny, suborders, and the problem of skeletal organization. In: Proceedings of the 11th International Bryozoology Association Conference (A. Herrera-Cubilla & J. B. C. Jackson, eds), pp. 87–103. Smithsonian Tropical Research Institute, Balboa, Republic of Panama.

Taylor, P. D. & Jones, C. G. (1993). Skeletal ultrastructure in the cyclostome bryozoan *Hornera*. Acta Zoologica, 74 (2), 135–143. https://doi.org/10.1111/j.1463-6395.1993.tb01230.x

Taylor, P. D. & Gordon, D. P. (2003). Endemic new cyclostome bryozoans from Spirits Bay, a New Zealand marine-biodiversity “hotspot”. New Zealand Journal of Marine and Freshwater Research. 37:653–669.

Taylor, P. D. & Waeschenbach, A. (2019). Phylogenetic affinities of *Crisulipora* and the dual origin of branch articulation in cyclostome bryozoan. Australasian Palaeontological Memoirs, 52: 155–161.

Taylor, P. D., Lombardi, C. & Cocito, S. (2015). Biomineralization in bryozoans: present, past and future. Biological Reviews, 90, 1118–1150. doi: 10.1111/brv.12148

Taylor, P. D., Waeschenbach, A., Smith, A. M., Gordon, D. P. (2015). In search of phylogenetic congruence between molecular and morphological data in bryozoans with extreme adult skeletal heteromorphy. Systematics and Biodiversity, 13: 525–544.

Taylor, P. D., Di Martino, E. & Martha, S. O. (2018). Colony growth strategies, dormancy and repair in some Late Cretaceous encrusting bryozoans: insights into the ecology of the Chalk seabed. Palaeobiodiversity and Palaeoenvironments 99, 1–22. https://doi.org/10.1007/s12549-018-0358-8.

Winston, J. E. (1978). Polypide morphology and feeding behavior in marine ectoprocts. Bulletin of Marine Science. 28(1):1–31.

Waeschenbach, A., Cox, C. J., Littlewood, D. T. J., Porter, J. S., Taylor, P. D. (2009). First molecular estimate of cyclostome bryozoan phylogeny confirms extensive homoplasy among skeletal characters used in traditional taxonomy. Molecular Phylogenetics and Evolution, 52: 241–251.

Waeschenbach, A., Taylor, P. D., & Littlewood, D. T. J. (2012). A molecular phylogeny of bryozoans. Molecular Phylogenetics and Evolution, 62, 718–735.

Wood, A. C. L., Probert, P. K., Rowden, A. A., Smith, A. M. (2012). Complex habitat generated by marine bryozoans: a review of its distribution, structure, diversity, threats, and conservation. Aquatic Conservation: Marine and Freshwater Ecosystems, 22: 547–563.

